# Conformational change of the Plasmodium TRAP I domain is essential for sporozoite migration and transmission of malaria

**DOI:** 10.1101/2022.08.24.505106

**Authors:** Friedrich Braumann, Dennis Klug, Jessica Kehrer, Chafen Lu, Timothy A. Springer, Friedrich Frischknecht

## Abstract

Eukaryotic cell adhesion and migration relies on surface adhesins connecting extracellular ligands to the intracellular actin cytoskeleton. *Plasmodium* sporozoites are transmitted by mosquitoes and rely on adhesion and gliding motility to colonize the salivary glands and to reach the liver after transmission. During gliding the essential sporozoite adhesin TRAP engages actin filaments in the cytoplasm of the parasite., while binding ligands on the substrate through its inserted (I)-domain. Crystal structures of TRAP from different *Plasmodium* species revealed the I-domain in closed and open conformations. Here, we probe the importance of these two conformational states by generating parasites expressing versions of TRAP with the I-domain stabilized in either the open or closed state with disulfide bonds. Strikingly, both mutations impact sporozoite gliding, mosquito salivary gland entry and transmission. Absence of gliding in sporozoites expressing the open TRAP I-domain could be partly rescued by adding a reducing agent. This suggests that dynamic conformational change is required for ligand binding, gliding motility and organ invasion and hence sporozoite transmission from mosquito to mammal.

## Introduction

Malaria-causing parasites undergo a complex life cycle between mosquitoes and vertebrate hosts. A key feature, essential during transmission to and from the mosquito, is parasite motility (Douglas et al., 2015). *Plasmodium* sporozoites develop in oocysts at the midgut wall and burst into the circulating fluid of the hemolymph followed by specific emigration into salivary glands (Frischknecht & Matuschewski, 2017). Within the salivary glands, sporozoites mature and gain the capacity to be highly motile once isolated or transmitted (Vanderberg, 1974). During the probing phase of a mosquito bite, the sporozoites are ejected with the saliva into the skin, where they migrate at high speed and enter into lymph and blood vessels (Hopp et al., 2021; Menard et al., 2013). Those entering the blood stream are transported throughout the circulatory system and arrest specifically in the liver, where they ultimately enter hepatocytes to differentiate intomerozoites, which subsequently infect red blood cells (Prudencio et al., 2006).

To navigate this complex journey (Figure 1A), sporozoites have a number of surface proteins including the single-pass transmembrane adhesin thrombospondin related anonymous protein (TRAP). TRAP likely evolved to bind multiple ligands found in the salivary gland, skin and liver (Klug et al., 2020; Matuschewski et al., 2002). TRAP harbors two extracellular domains, the I-domain and the thrombospondin type I repeat, TSR. On the cytoplasmic side, a small tail links to actin filaments that are formed underneath the plasma membrane (Heintzelman, 2015) (Figure 1B). The I-domain is found in many other proteins, including integrins of multicellular organisms. Deletion of *trap* allows sporozoite formation and exit from the oocysts but *trap(−)* parasites largely fail to enter the salivary glands (Sultan et al., 1997). Hemolymph-derived *trap(−)* sporozoites also show defects in substrate adhesion and fail to undergo typical gliding motility (Figure 1C) and instead can only move back and forth over a single adhesion site, a phenomenon termed patch gliding (Münter et al., 2009). Deletion of the I-domain largely phenocopies deletion of the entire protein and mutations of or around the metal ion dependent adhesion site (MIDAS) compromise sporozoite infectivity (Klug et al., 2020; Matuschewski et al., 2002).

**Figure 1.**
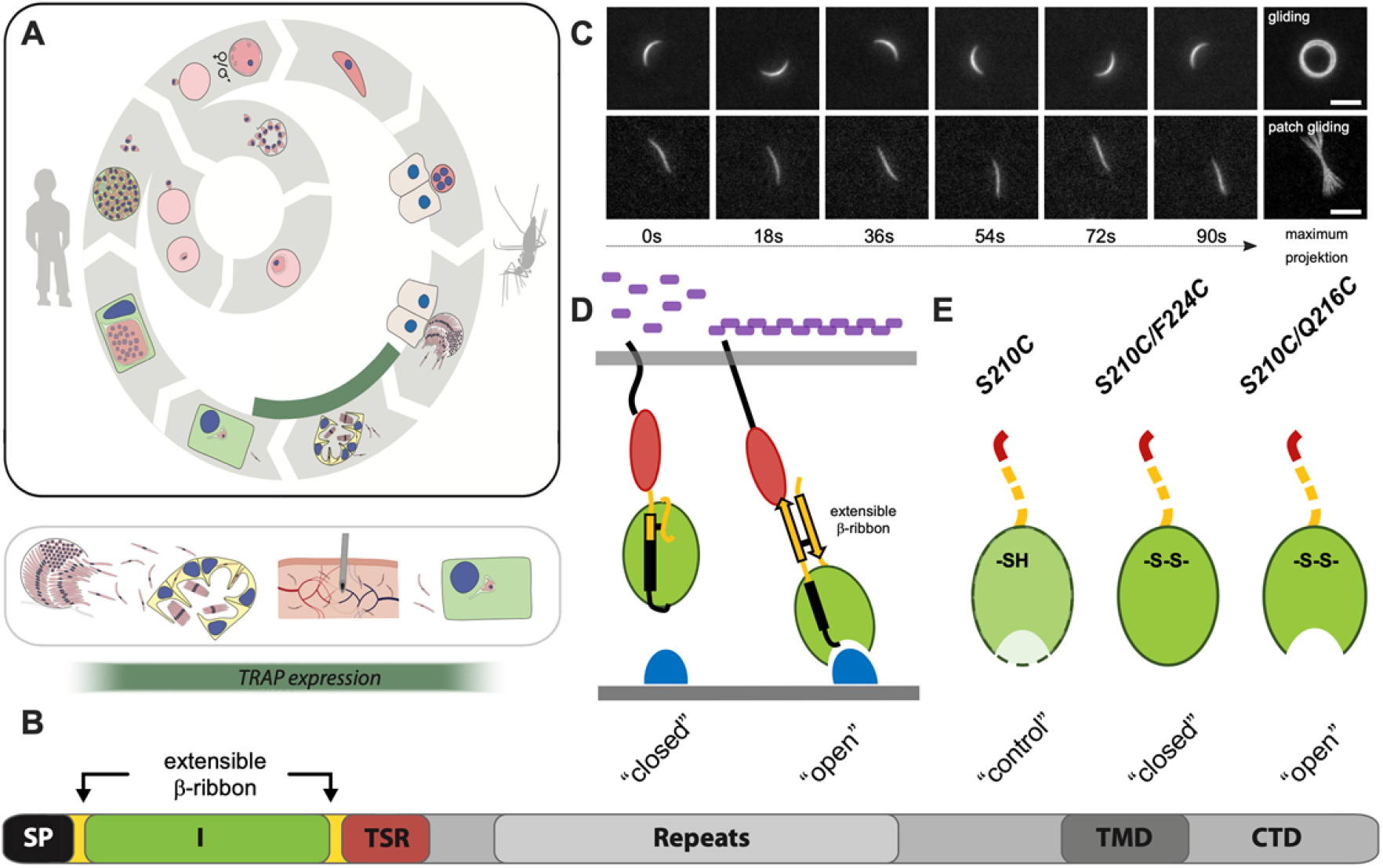
TRAP mediated sporozoite migration and potential role for I domain. **A**: A simplified schematic overview of the *Plasmodium* life cycle. The part of the life cycle in which TRAP is expressed is highlighted by a green bar. Note that sporozoites have to actively enter salivary glands, cross through the dermis and enter hepatocytes. **B**: Domain architecture of full-length TRAP showing signal peptide (SP), I domain (I, green), thrombospondin type I repeat (TSR), repeats, transmembrane domain (TMD), and cytoplasmic tail domain (CTD). **C**: Movie stills (first 6 panel in each row) and maximum intensity projections (7th panel in each row) of a 90 second video of *P. berghei* sporozoites undergoing normal circular gliding motility (top row) and patch gliding (bottom row). Scale bars: 5 μm. **D**: Illustration of TRAP linkage to the cytoskeleton and ligand binding. TRAP I domain in the “closed” conformation is unbound and has low adhesion and no signaling activity. TRAP I domain in the “open” conformation binds to substrates potentially leading to intracellular signaling and interaction with actin filaments. **E**: Cartoon of the three different mutants generated in this study. and *cmTRAP∷S210C/F224C* fixes the I domain in the closed state and *cmTRAP*∷*S210C/Q216C* fixes the I domain in the open state. The *cmTRAP∷S210C* mutant containing only one cysteine serves as control parasite line.

Crystal structures of the N-terminal portion of TRAP containing the TSR and I-domain in human-infective *Plasmodium* spp. revealed the I domain in both open and closed conformations ((Song et al., 2012) Figure 1D). Analogy to other I domains Shimaoka et al., 2002 suggests that the open state of TRAP has higher ligand-binding affinity due to the changed position of a ligated Mg^2+^ ion within the MIDAS, which also impacts the conformation of the entire N-terminal domain, such that in the open state an extensible β-ribbon appears between the I and TSR domains (Shimaoka et al., 2002; Song et al., 2012). Binding to ligand therefore stabilizes the open conformation. Force applied by the actin cytoskeleton to TRAP that is resisted by extracellular ligand would transmit tensile force through TRAP, and further stabilize its extended, open conformation with the extensible β-ribbon. A similar force-dependent process appears to stabilize the extended, open conformation of integrins (Li & Springer, 2017). Hence, ligand binding, together with tensile force, likely stabilizes a strong conformational change in TRAP that activates gliding motility. In integrins, stabilizing the domain in an open state by engineered disulphide bonds increased ligand affinity ~1,000-fold, while stabilizing it in a closed state gave wild type-like affinities (Shimaoka et al., 2001).

To investigate if the two I domain conformational states are also important for sporozoite migration and infectivity, we designed closed and open versions of the *Plasmodium berghei* TRAP I domain through addition of two cysteines at different positions. Both mutations impacted the capacity of sporozoites to migrate, enter salivary glands and infect mice. Interestingly, hemolymph-derived sporozoites expressing the open conformation could not migrate on flat surfaces but still entered into salivary glands, albeit at a reduced level compared to wild type. Sporozoites derived from the salivary gland remained immotile but their motility could be partly rescued by the addition of the reducing agent dithiothreitol (DTT).

## Results

### *P. berghei* sporozoites expressing TRAP with conformationally stabilized I domains fail to transmit

Previous work on conformation stabilizing I domain mutations in integrins (Shimaoka et al., 2001; Shimaoka et al., 2003) and the crystal structures of *Plasmodium* I domains (Song et al., 2012) informed on the positions to introduce cysteines for engineering disulphide bonds. To generate TRAP mutants with an I domain stabilized in the closed state, we substituted cysteines for Ser-210 and Phe-224 in a codon modified version of *trap*. Similarly, we mutated Ser-210 and Gln-216 to cysteine to stabilize the I domain in the open state, and used a single S210C mutant as a control (**Figure 1E** and **Figure S1**). These TRAP constructs were ligated into a previous vector (Klug et al., 2020) in order to replace the endogenous TRAP with the three different constructs (**Figure S1**). Constructs were either transfected into the fluorescent line *cFluo*, expressing EGFP constitutively from the *ef1α* promoter and mCherry in insect stages from the *csp* promoter, or a non-fluorescent *trap(−)* line (Bane et al., 2016; Klug et al., 2020). Generation of clonal lines yielded the parasite lines *S210C* (control, fluorescent), *S210C/F224C* (closed, non-fluorescent) and *S210C/Q216C* (open, fluorescent). For all experiments the *cFluo* line was used as the control and will be referred to as wild type in the following. As expected, all lines showed no growth difference to wild type in mice and a similar range of mosquito infection with the numbers of oocysts even being increased for the closed and open mutants (**Figure 2A**). Sporozoites also egressed efficiently from oocysts and could be isolated from the hemolymph. Western blotting of hemolymph derived sporozoites showed similar expression of TRAP in the wild type, control and conformationally stabilized parasite lines (**Figure 2B**). Immunofluorescence analysis of hemolymph derived sporozoites that were fixed but not permeabilized revealed similar signals of TRAP on the surface in all lines (**Figure 2C**). The numbers of sporozoites were similar in the midgut and hemolymph for all lines (**Figure 3A,C, Table 4**) but showed a dramatic drop in salivary gland residency of the lines expressing TRAP in the closed and open conformations (**Figure 3B,D, Table 4**).

**Figure 2.**
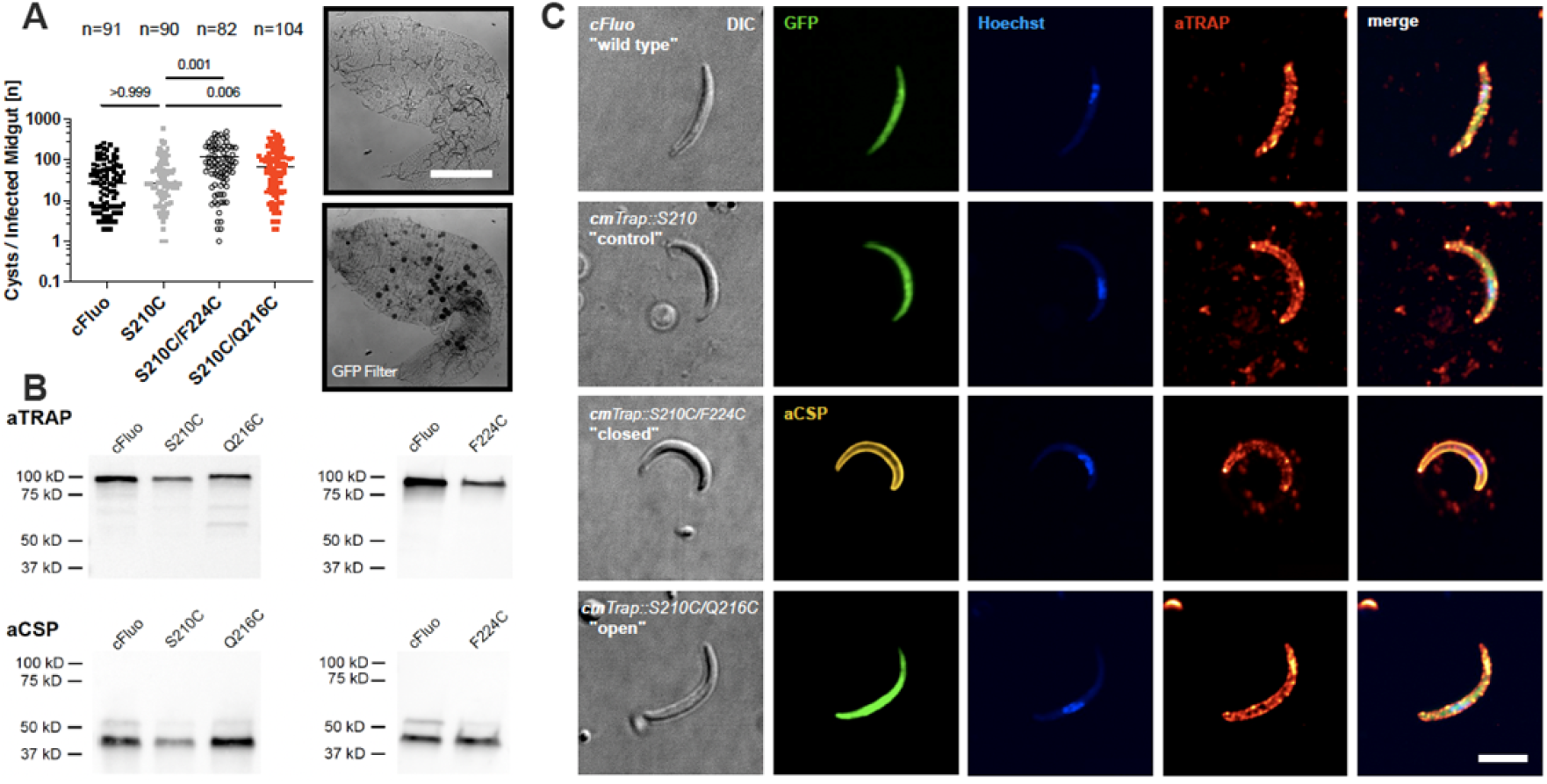
Modified TRAP proteins are expressed and localize as in wild-type controls. **A**: Numbers of oocysts in infected midguts of the different lines. Values (n) above dot plots indicate numbers of observed midguts from at least three independent cage feeds and numbers above horizontal bars indicate p-values. Data was tested for significance using the Kruskal-Wallis-test. Images show *cFluo* oocysts in an infected midgut without (top) and with (bottom) green fluorescence. Scale bar: 200 μm. **B**: Western blots using an anti-TRAP antibody of the control lines *cFluo* and *S210C* in comparison to *S210C/Q216C* (open) and *S210C/F224C* (closed). Anti-CSP antibody mAb 3D11 was used as loading control, revealing the two expected CSP-bands (Coppi et al., 2011). **C**: Surface expression of TRAP mutants on fixed but not permeabilized sporozoites. Differential interference contrast (DIC) and immunofluorescence images of representative examples of hemolymph derived sporozoites of the indicated parasite lines stained with anti-TRAP antibodies shown in red (Klug et al., 2020). Hoechst was used to reveal the sporozoite nuclear DNA shown in blue. Cytoplasmic GFP or staining with anti-CSP antibodies were used to validate the integrity of sporozoites. Scale bar: 5 μm.

**Figure 3.**
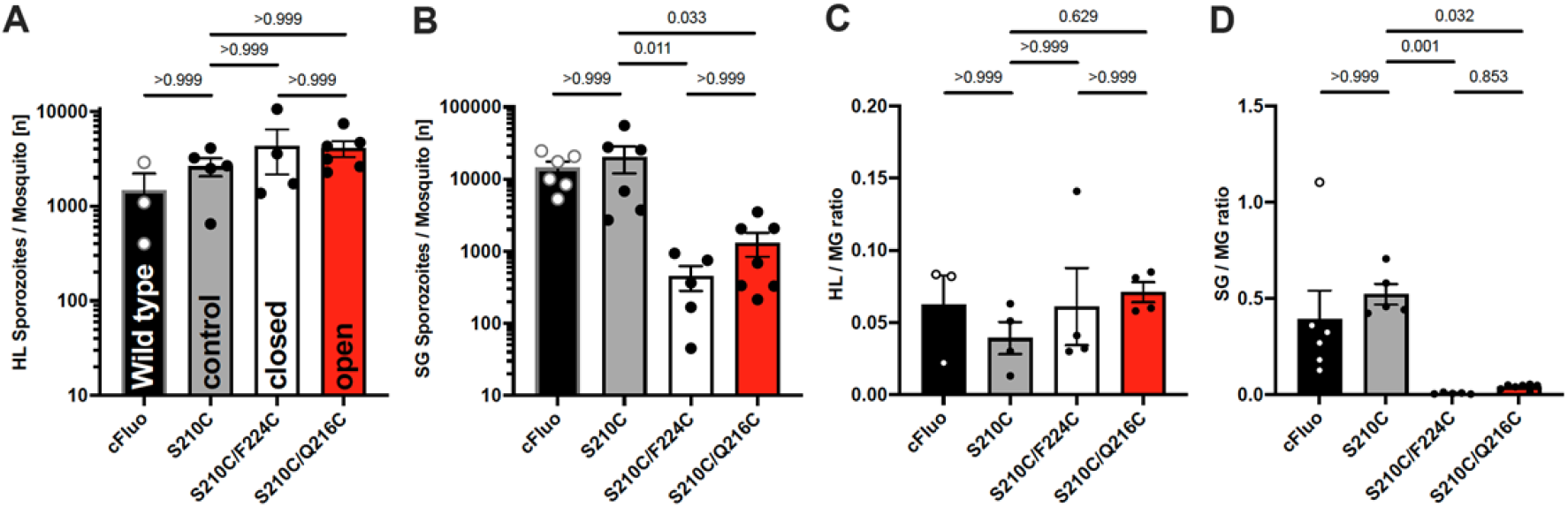
Sporozoites with fixed I-domains fail to invade salivary glands. **A**: Numbers of hemolymph (HL) derived sporozoites reveal no difference between the parasite lines. **B**: Numbers of salivary gland derived (SG) sporozoites reveal strong reduction in salivary gland invasion for the closed and open I domain expressing parasite lines. **C**: Ratios of hemolymph-derived versus oocyst-derived sporozoites reveal no difference between the parasite lines. **D**: Ratios of salivary gland derived versus hemolymph derived sporozoites reveal strong reduction in salivary gland invasion for the closed and open I domain expressing parasite lines. Data shown in (**A-D**) represent pooled results from 3-7 independent experiments. Shown is mean +/− SEM. Dots represent data from individual mosquito feedings. Data was tested for significance with the Kruskal-Wallis-test.

To investigate the capacity to infect mice, we isolated sporozoites of all three lines as well as the wild type from the hemolymph and injected 10,000 sporozoites of each line into individual naïve C57Bl/6 mice. This showed the expected 100% infection rate in wild type and S210C control parasites with a prepatent period (time until parasites can be detected in red blood cells) of around 5 days (**Figure 4A-C** and **Table 1**). However, hemolymph sporozoites expressing the closed or open TRAP I domains infected only around 50% of mice and those infected showed a prolonged prepatent period of 6-7 days (**Figure 4A-C** and **Table 1**). We next determined how the decreased colonization of the salivary glands by the closed and open mutants affected transmission of sporozoites. To this end we allowed ten infected mosquitoes to bite a naïve mouse and determined the onset and development of a blood stage infection. As expected, the two control lines (wild type and S210C) infected all mice and showed prepatent periods of 3-4 days (**Figure 4D-F** and **Table 1**). In contrast, mosquitoes that were infected by the closed mutant (S210C/F224C) could not infect mice while one out of ten mice bitten by mosquitoes harboring the open mutant (S210C/Q216C) was infected (**Figure 4D-F** and **Table 1**). This mouse showed a delayed onset of blood stage infection at 6 days post infection. The number of salivary gland-derived sporozoites in the open mutant (S210C/Q216C) was higher, but not significantly higher, than in the closed mutant (S210C/F224C) (**Figu,re 3C)**; however, we never managed to isolate sufficient numbers of sporozoites from salivary glands of the closed mutant (S210C/F224C) to conduct further experiments. We therefore isolated only sporozoites from salivary glands of mosquitoes infected by the control lines or the open mutant (S210C/Q216C) and injected these into naïve mice. This showed that three out of five mice infected with S210C/Q216C salivary gland sporozoites became blood stage patent, albeit again with a 2-day delay compared to the controls, while all mice infected with the controls became blood stage positive (**Figure 4G-I** and **Table 1**). This suggests that both mutants are severely impacted in their capacity to transmit from mosquitoes to mammals (**Table 4**).

**Figure 4.**
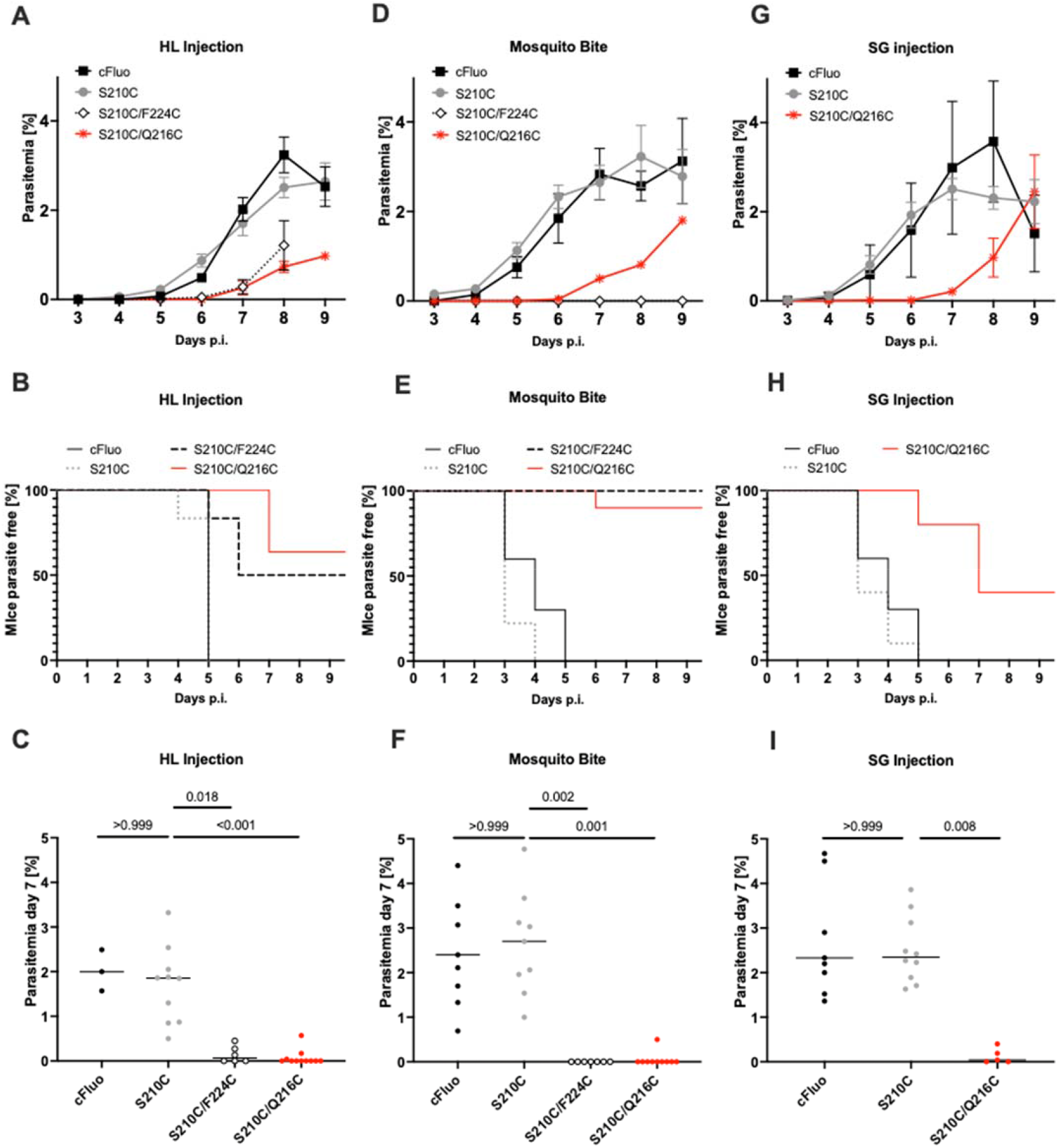
Conformational stabilization of the TRAP I domain impedes transmission. **A**: Blood stage parasitemia after injection of 10,000 hemolymph-derived (HL) sporozoites of the different lines into C57Bl/6 mice from day 3 to day 9 post infection. Shown is the mean +/− SEM. Only data of infected mice has been plotted. **B**: Kapplan Meier plot showing the percentage of parasite free mice until day 9 of infection. **C**: Parasitemia on day 7 post hemolymph sporozoite (HLS) injection, tested for significance with Kruskal Wallis test. **D**: Blood stage parasitemia after C57Bl/6 mice were bitten by mosquitoes (ten mosquitoes/mouse) infected with the indicated parasite lines at day 0. Shown is the mean +/− SEM where appropriate. Only data of infected mice has been plotted for all but the S210C/F224C (closed I domain) infections, where all mice stayed negative. **E**: Kapplan Meier plot showing the percentage of parasite free mice after mosquito bites until day 9 of infection. **F**: Parasitemia on day 7 post mosquito bite, tested for significance with Kruskal Wallis test. **G**: Blood stage parasitemia after injection of 10.000 salivary gland-derived sporozoites of the indicated parasite lines into C57Bl/6 mice at day 0. Only data of infected mice has been plotted. **H**: Kapplan Meier plot showing the percentage of parasite free mice after salivary gland sporozoite injection until day 9 of infection. **I**: Parasitemia on day 7 post salivary glands sporozoite injection, tested for significance with Kruskal-Wallis-test.

**Table 1.** Transmission potential of the generated parasite lines *S210C*, *S210C/Q216C* and *S210C/F224C* in comparison to the control *cFluo*. The prepatency is determined as the time between infection and the first observance of blood stages and is given as the mean of all mice that became blood stage positive; n.d.: not determined, n.a.: not applicable.

| Parasite line | Inoculation type | Mice infected / total exposed | Prepatency (days) |
| --- | --- | --- | --- |
| **cFluo** | HL iv | 3/3 | 5.0 |
|  | SG iv | 10/10 | 3.9 |
|  | Mosquito Bite | 10/10 | 3.9 |
| **S210C** | HL iv | 10/10 | 4.8 |
|  | SG iv | 10/10 | 3.5 |
|  | Mosquito Bite | 9/9 | 3.2 |
| **F224C** | HL iv | 3/6 | 5.7 |
|  | SG iv | n.d. | n.d. |
|  | Mosquito Bite | 0/10 | n.a. |
| **Q216C** | HL iv | 3/11 | 7.0 |
|  | SG iv | 3/5 | 6.3 |
|  | Mosquito Bite | 1/10 | 6.0 |

### Conformationally stabilized open mutants do not glide

As gliding motility is a key factor in sporozoite infectivity, we next investigated the capacity of the control and mutant lines to glide in a standard gliding assay. To this end, sporozoites were either isolated from the hemolymph or salivary glands (if possible) and placed in RPMI medium containing 3% bovine serum albumin (BSA), which stimulates motility (Vanderberg, 1974). We scored the different patterns of motility that could be observed (**Figure S2**). This analysis showed robust gliding motility for the control parasites isolated from both hemolymph or salivary glands, which was comparable to wild type (**Figure 5A,B**). In contrast, hemolymph derived sporozoites expressing TRAP in either the closed or open form, showed strongly impaired gliding motility. Sporozoites expressing the closed I domain (S210C/F224C) showed low levels of persistent gliding (~10% of control level). Strikingly, despite the observation of hundreds of sporozoites expressing the open I domain (S210C/Q216C), none of these were gliding (**Figure 5A**). This observation is somewhat counterintuitive, since sporozoites expressing the open variant (S210C/Q216C) showed some level of colonization of the salivary glands (**Figure 3A-D**). Isolation of sporozoites expressing the open I domain (S210C/Q216C) from the salivary glands also showed no motility, while the controls moved as expected (**Figure 5B**). Interestingly, salivary gland sporozoites expressing the open I domain (S210C/Q216C) showed nearly no floating parasites compared to the controls S210C and *cFluo* (**Figure 5B**) suggesting enhanced adhesion. Intriguingly, hemolymph derived sporozoites expressing the closed (S210C/F224C) were significantly faster during migration than corresponding controls (**Figure 5C**). This hints to a decrease in adhesion capacity of the closed mutant, which initially speeds up parasites but also reduces their capacity to stay attached to the substrate, as observed before for sporozoites treated with drugs that affect actin polymerization (Münter et al., 2009).

**Figure 5.**
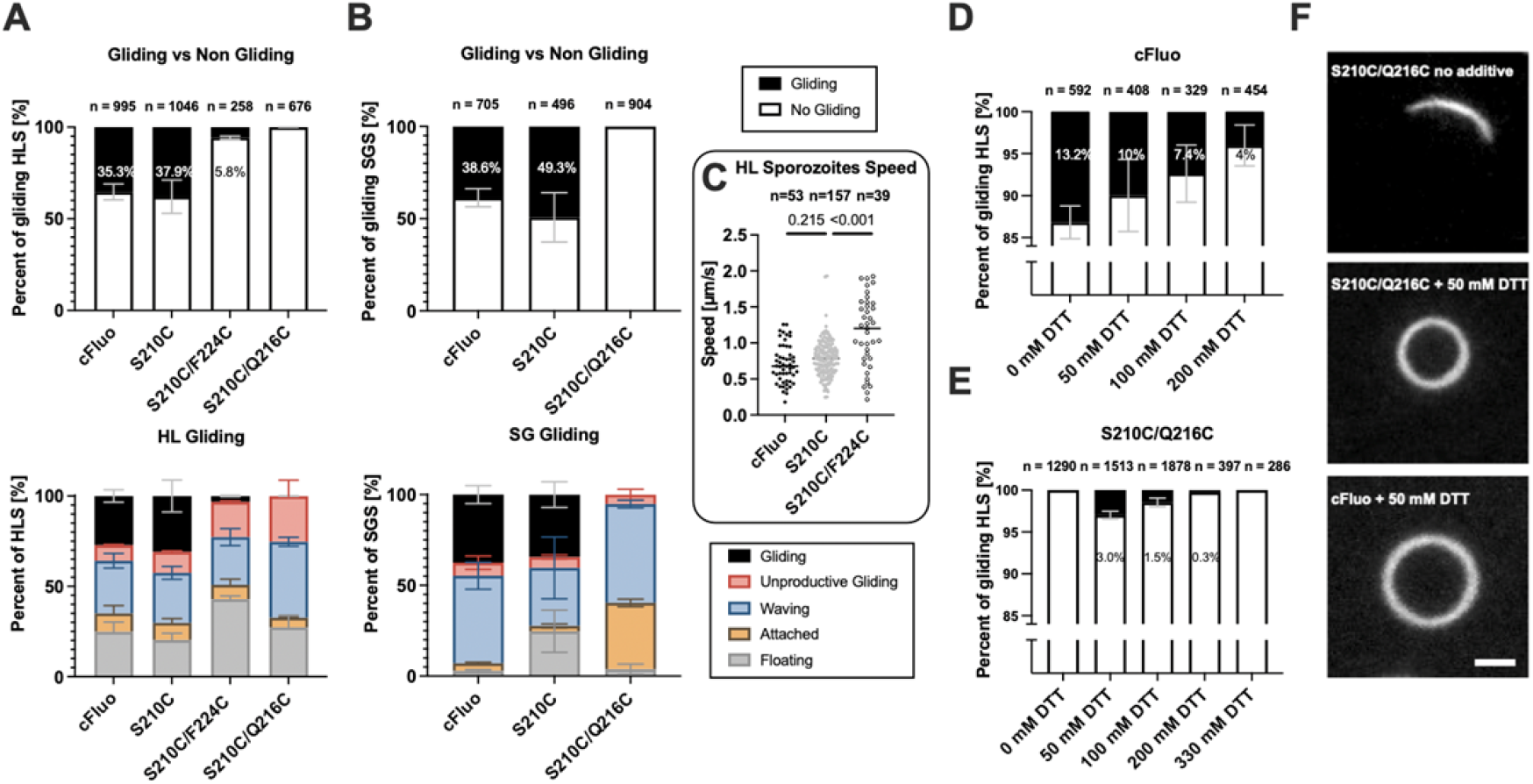
Sporozoites expressing the open TRAP I domain do not glide unless treated with DTT. **A**: Movement type analysis of hemolymph derived sporozoites (HLS) of the indicated parasite lines. Top: Classification of attached sporozoites by motility. Bottom: Classification by motility type. n indicates the number of observed sporozoites. See Figure S2 for the classification of different movement patterns. Floaters have been omitted in the analysis presented in the top graph. **B**: Movement type analysis of salivary gland derived sporozoites (SGS) of the indicated parasite lines. n indicates the number of observed sporozoites. See Figure S2 for the classification of different movement patterns. Floaters have been omitted in the analysis presented in the top graph. **C**: Speed of hemolymph derived productively gliding sporozoites from the indicated lines. Data were tested for significance with Kruskal-Wallis-test. **D**: Percentage of gliding and non-gliding control hemolymph sporozoites at increasing concentrations of DTT. Data were generated from at least two independent mosquito feedings. n indicates the number of observed sporozoites. Floaters are included in the non-motile fraction. **E**: Percentage of gliding hemolymph sporozoites expressing the open TRAP I domain at increasing concentrations of DTT. Data represent pooled data from at least two independent mosquito feedings. n indicates the total number of observed sporozoites. Floaters are included in the non-motile fraction. **F**: Maximum projections (1 min time lapse, 3 s per frame) of single *S210C/Q216C* sporozoites in presence or absence of 50 mM DTT. A projection from a motile sporozoite of the *cFluo* line treated with 50 mM DTT is shown as control. Scale bar: 5 μm.

### A reducing agent rescues motility in TRAP open I domain expressing sporozoites

Disulphide bonds can be opened by a reducing agent. Hence, we hypothesized that adding reducing agents to the medium might reconstitute motility of sporozoites expressing the open I domain (S210C/Q216C). However, there are other structurally important disulphide bonds present in TRAP as well as in other proteins on the sporozoite surface including the circumsporozoite protein (CSP), the major surface protein of sporozoites (Doud et al., 2012). Nevertheless, we aimed to find a condition, in which the advantage gained by disrupting our introduced disulphide bonds outweighs the disruption of other disulphide bonds. To increase the likelihood of finding an ideal experimental setup, we tested two reducing agents, dithiothreitol (DTT) and tris(2-carboxyethyl)phosphine (TCEP), which differ in their ability to cross membranes and, because of structural differences, in their potential to reduce disulfide bonds in folded proteins (Cline et al., 2004). We screened a range of DTT and TCEP concentrations and incubation times. By doing so we found that gliding assays could not be performed in the simultaneous presence of BSA and reducing agents, as amorphous aggregates formed (Yang et al., 2015). Instead, we incubated sporozoites with DTT or TCEP before starting the actual assay. The reducing agents were then washed out and the sporozoites were activated with RPMI medium supplemented with 3% BSA. Notably, this prolonged protocol resulted in a reduction of approximately 60% in productively moving wild-type sporozoites which has to be taken into account when comparing results from regular gliding assays with assays performed with DTT and TCEP treated sporozoites (**Figure 5D**). We first determined at which concentration of DTT or TCEP, wild type sporozoites were still motile (**Table 2**). This showed that already after 15 min incubation, a significant reduction of motility could be observed at 50 mM of DTT and at 5 mM of TCEP. At concentrations of 300 mM DTT and 50 mM TCEP respectively, no motility was observed anymore (**Figure 5D**, **Figure S3**, **Table 2**). We therefore incubated hemolymph derived sporozoites expressing the open I domain (S210C/Q216C) in different low concentrations of DTT and TCEP. Gliding assays under reducing conditions were only performed with sporozoites expressing TRAP fixed in the open state (S210C/Q216C) as sporozoites of this mutant showed no productive gliding under regular conditions. Gliding assays with reducing agents revealed conditions in which DTT could rescue motility of sporozoites expressing the open I domain (S210C/Q216C) (**Table 3**). At 50 and 100 mM 3% and 1.5% of sporozoites were found productively moving, respectively (**Figure 5E**, **Figure S3**). This compares to 13% of control parasites moving in the absence of DTT and 10% and 7.4% moving at 50 and 100 mM DTT, respectively (**Figure 5D**). Hence, using these conditions motility was rescued to approximately 20-30% of wild type level. These data suggest that DTT can reduce the introduced disulphide bond and enable some TRAP molecules on the sporozoite surface to function normally leading to an increase of migrating sporozoites.

**Table 2.**
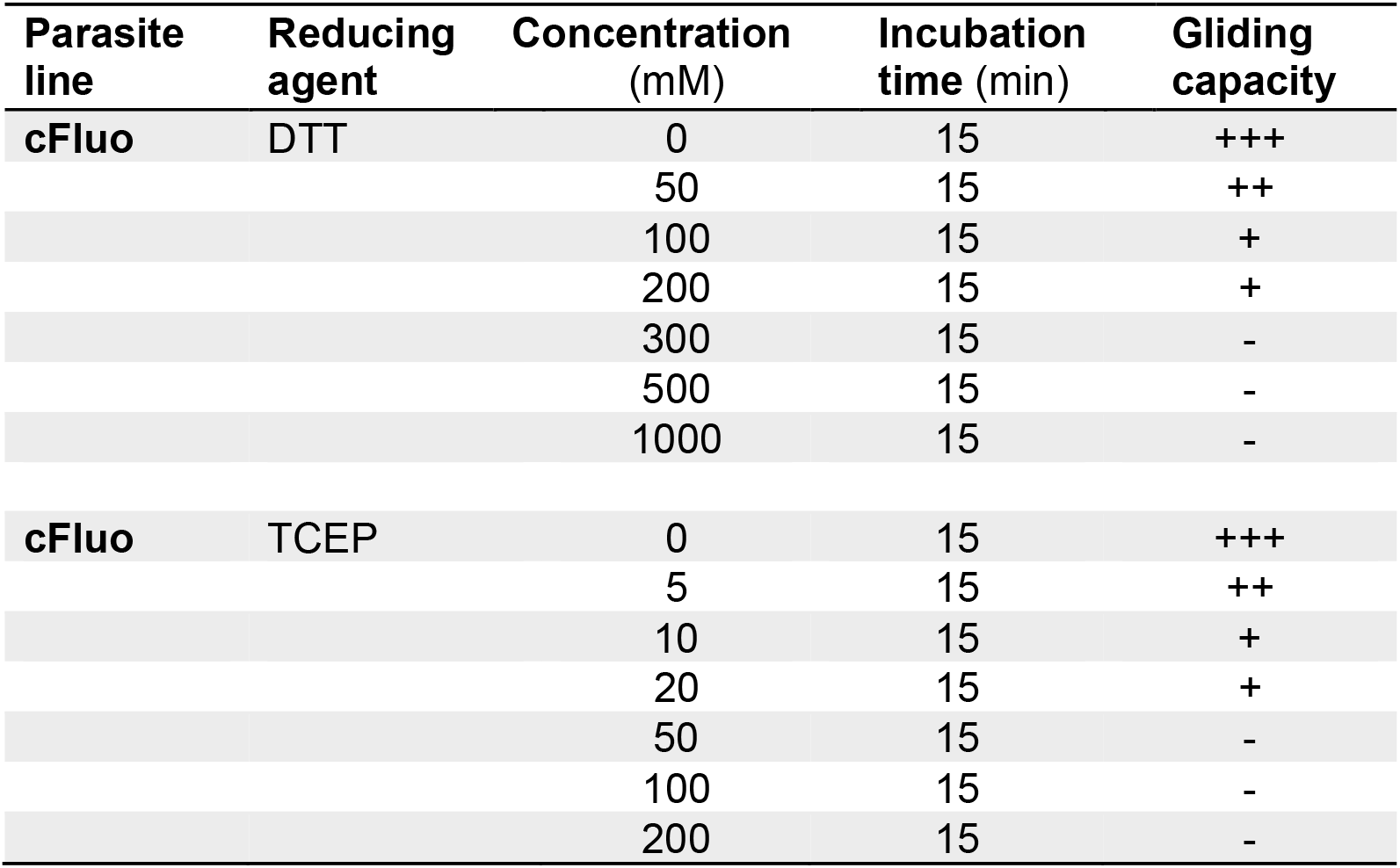
Screening for TCEP and DTT concentrations affecting wild-type motility.

| Parasite line | Reducing agent | Concentration (mM) | Incubation time (min) | Gliding capacity |
| --- | --- | --- | --- | --- |
| cFluo | DTT | 0 | 15 | +++ |
|  |  | 50 | 15 | ++ |
|  |  | 100 | 15 | + |
|  |  | 200 | 15 | + |
|  |  | 300 | 15 | - |
|  |  | 500 | 15 | - |
|  |  | 1000 | 15 | - |
| cFluo | TCEP | 0 | 15 | +++ |
|  |  | 5 | 15 | ++ |
|  |  | 10 | 15 | + |
|  |  | 20 | 15 | + |
|  |  | 50 | 15 | - |
|  |  | 100 | 15 | - |
|  |  | 200 | 15 | - |

**Table 3.** Incubation of open I domain expressing sporozoites (*S210C/Q216C*) with DTT but not TCEP rescues motility.

| Parasite Line | Reducing Agent | Concentration (mM) | Incubation time (min) | Gliding capacity |
| --- | --- | --- | --- | --- |
| S210C/Q216C | DTT | 5 | 15 | - |
|  |  | 30 | 2 | - |
|  |  | 30 | 15 | - |
|  |  | 50 | 15 | + |
|  |  | 100 | 2 | - |
|  |  | 100 | 15 | + |
|  |  | 100 | 45 | - |
|  |  | 200 | 15 | - |
|  |  | 330 | 15 | - |
| S210C/Q216C | TCEP | 1 | 15 | - |
|  |  | 2 | 60 | - |
|  |  | 5 | 15 | - |
|  |  | 10 | 2 | - |
|  |  | 10 | 15 | - |
|  |  | 20 | 15 | - |
|  |  | 100 | 15 | - |

**Table 4.** Absolute sporozoite numbers in midgut (MG), hemolymph (HL) and salivary glands (SG) of all analyzed parasite strains. Last column shows countings from infected mosquito cages that revealed over 1000 sporozoites per salivary gland (on average) from at least 10 dissected mosquitoes. Shown is mean ± SD. Bold numbers indicate significant difference from controls.

|  | No. of HL<br>Sporozoites | No. of MG<br>Sporozoites | No. of SG<br>Sporozoites | HL/MG<br>Ratio | SG/MG<br>Ratio | Mosquitoes<br>with > 1000 spz |
| --- | --- | --- | --- | --- | --- | --- |
| **cFluo** | 1.500±1.300 | 28.000±20.000 | 14.000±7.600 | 0.06±0.04 | 0.39±0.36 | 6/6 |
| **S210C** | 2.600±1.300 | 57.000±22.000 | 20.000±20.000 | 0.04±0.03 | 0.54±0.13 | 6/6 |
| **S210C/Q216C** | 4.100±1.700 | 52.000±40.000 | **1.300±1.300** | 0.07±0.01 | **0.04±0.01** | **3/7** |
| **S210C/F224C** | 4.300±4.300 | 73.000±32.000 | **450±380** | 0.06±0.05 | **0.01±0.01** | **0/5** |

## Discussion

Here we report evidence that a conformational change in the I domain of the *Plasmodium* sporozoite surface protein TRAP is important for gliding motility and parasite transmission. The transition from a closed to an open state was suggested to be important based (i) on structural studies that revealed the I domain of *P. falciparum* and *P. vivax* TRAP in the closed and open conformations, respectively (Song et al., 2012) and (ii) analogy to the mechanism of action of I domains in human integrins (Shimaoka et al., 2003). In a closed state the integrin I domain has a low affinity for ligand binding, which is greatly increased in the open state. Thus, a ligand bound ‘open’ TRAP might signal back to the cytoplasm to change the state of actin filament assembly or TRAP association with actin filaments (Quadt et al., 2016; Song et al., 2012). By fixing the closed or open state with disulphide bonds, we found that both parasite lines expressing these variants are severely compromised in their capacity to transmit from mosquitoes to mice. This phenotype indicates that a key feature of TRAP function during gliding and invasion is the dynamic interchange between a closed and an open conformation of the I domain.

Conformational changes have also been reported for the major surface protein, the GPI-anchored circumsporozoite protein CSP (Coppi et al., 2011; Herrera et al., 2015). Here, the N-terminus appears to shield the single TSR of CSP and hence avoids critical ligand binding via the TSR until necessary. Yet, in CSP the N-terminus is also cleaved off by a protease and hence the protein might not undergo conformational change *per se*. Similarly, TRAP cleavage by rhomboid proteases also occurs and is important for gliding (Ejigiri et al., 2012). Several proteins have been suggested to specifically bind to TRAP: fetuin-A, a hepatocyte-specific protein (Jethwaney et al., 2005), saglin, an *Anopheles* salivary gland protein (Ghosh et al., 2009), alpha-v-containing integrins (Dundas et al., 2018) and platelet derived growth factor beta (Steel et al., 2021). We have previously called into question the specificity of these interactions by reporting that an I domain from *Toxoplasma gondii* MIC2, and to a lesser extent from the human integrin αXβ2, can functionally replace the I domain of *Plasmodium berghei* TRAP (Klug et al., 2020). In analogy to certain human I-domains such as αXβ2 and αMβ2 with promiscuous ligand binding (Springer, 2006), we interpreted these results such that TRAP evolved to bind multiple ligands. Our current data cannot resolve this controversy but suggest that screening for ligand binding might benefit from TRAP domains that are fixed in the open or closed state.

Antibodies against sporozoite surface proteins, especially CSP, are being explored for passive immunization and to inform the improvement of CSP-based vaccines (Aliprandini et al., 2018; Flores-Garcia et al., 2021; Gaudinski et al., 2021; Julien & Wardemann, 2019; Murugan et al., 2020; Wang et al., 2021). CSP is the predominant antigen on the sporozoite surface with TRAP being among the most abundant other surface proteins. Yet, compared to CSP, most of the TRAP is stored in secretory micronemes and only a small proportion is exposed on the sporozoite surface (Carey et al., 2014; Kehrer et al., 2016). Antibodies against TRAP have been generated and probed for their capacity to inhibit sporozoite gliding. While a first report of such inhibitory capacity (Spaccapelo et al., 1997) could not be reproduced (Gantt et al., 2000), several studies explored TRAP as a vaccine candidate (Kim et al., 2020; Lu et al., 2020; Nazeri et al., 2020; Tiono et al., 2018). Our work suggests that antibodies that stabilize the conformation of the TRAP I domain in either the open or the closed state could stop parasite migration in the skin or during liver entry. Gliding motility is affected differently in the parasites expressing TRAP with the I domain fixed in the two states. The closed mutant (S210C/F224C) shows faster motility likely because of weakened adhesion capability (**Figure 5C**). The open mutant (S210C/Q216C) appears to adhere normally or even enhanced to the substrate yet does not move productively, suggesting that it cannot detach from the substrate. This could however also be based on possible differences in the trapping efficiency of the introduced disulphide bonds leading to a stronger stabilization of the open conformation in the S210C/Q216C mutant compared to the closed state in the S210C/F224C mutant. As a consequence, some TRAP mutants might be partially able to transition between the open and closed states, enabling some movement, including faster movement by the mutant stabilized in the closed state. Still, the difference between the two mutants in their capability to move productively as well as the strong adhesion of salivary gland sporozoites expressing the open I domain (S210C/Q216C) indicates a difference in the ligand binding affinity as reported for human integrins with an I domain trapped in the open state (Shimaoka et al., 2003). This is also reminiscent of a mutant where TRAP cannot be cleaved by rhomboid proteases (Ejigiri et al., 2012). Rhomboid proteases cleave TRAP and hence ensure detachment from the substrate for efficient gliding motility. Mutations in TRAP that prevented cleavage by rhomboid proteases yielded sporozoites with severely compromised motility that struggled to enter salivary glands. The inability of sporozoites with the TRAP I domain to migrate suggests that sporozoites rely on both, a dynamic conformational change in the I domain and rhomboid mediated cleavage to ensure rapid gliding motility, which proceeds at a rate 10 times faster than rapidly migrating human cells such as neutrophils.

In conclusion, we suggest that the capacity of the TRAP I domain to dynamically switch between two conformational states is essential for *Plasmodium* sporozoite migration and transmission. Anti-TRAP antibodies that fix the I domain in one of the conformational states could constitute new tools to prevent malaria infection alone or in combination with anti-CSP antibodies.

## Acknowledgements

We thank Miriam Reinig and other members of the Frischknecht lab for mosquito rearing and Franziska Hentzschel for comments on the manuscript.

---

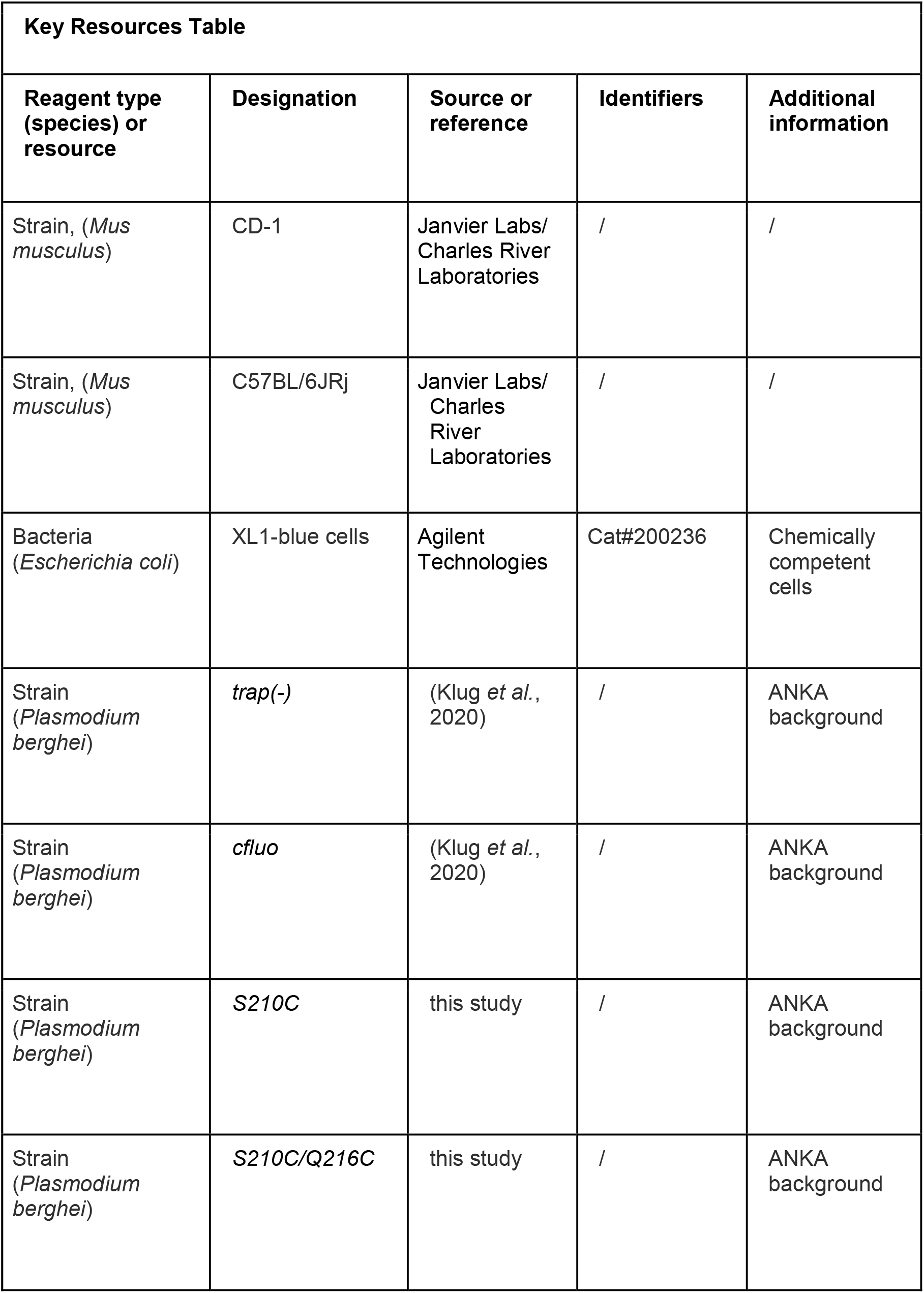
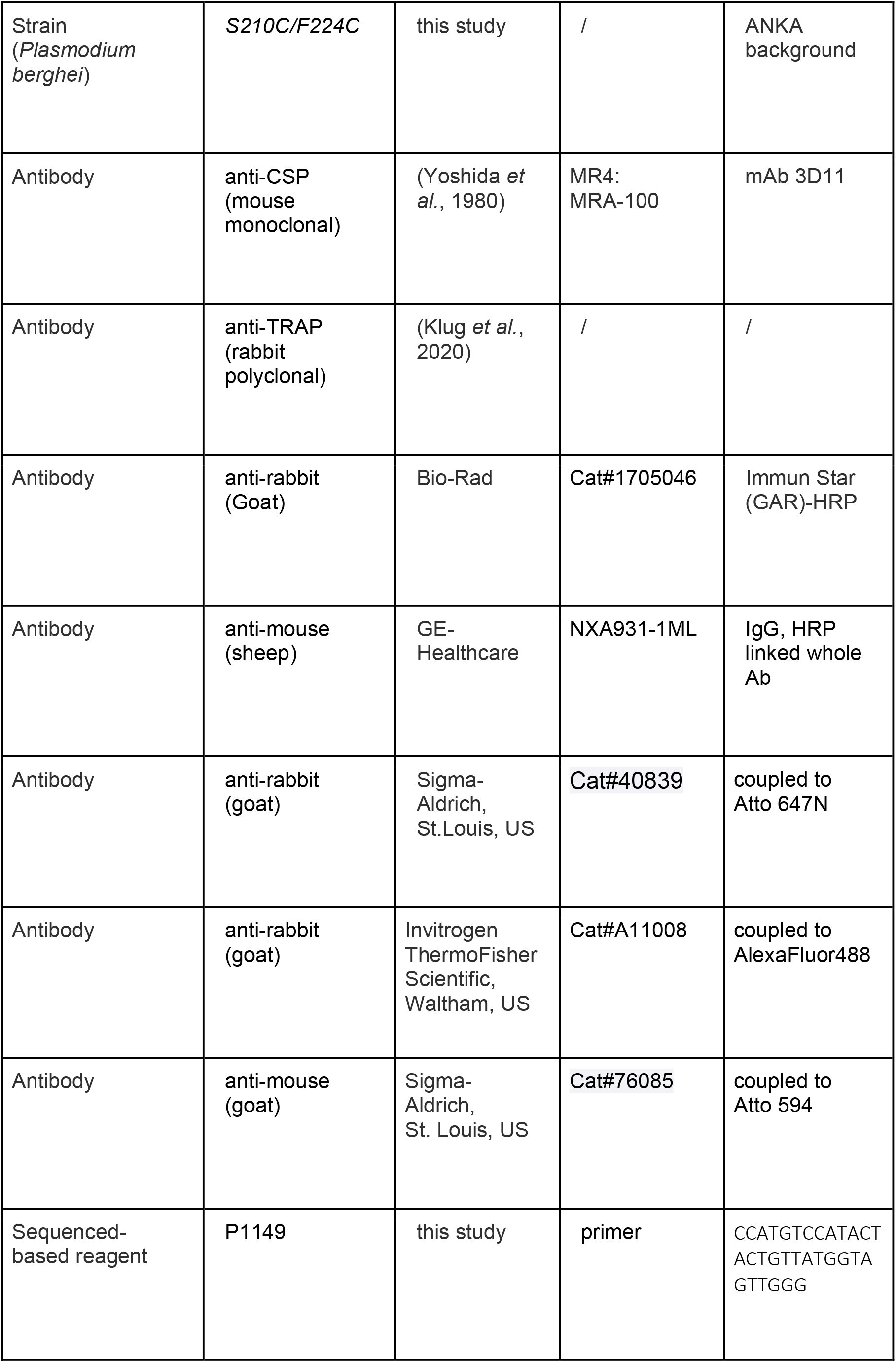
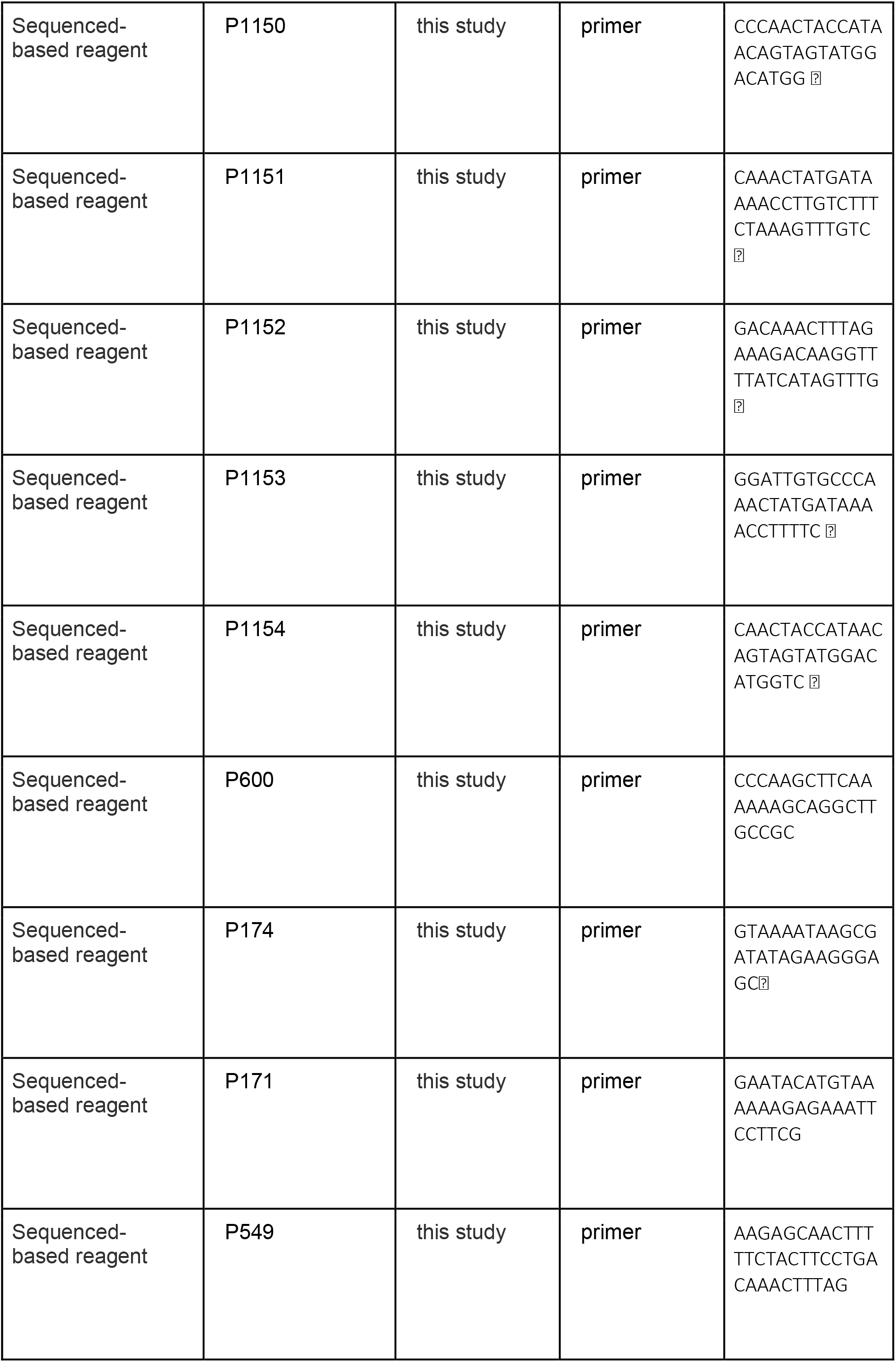
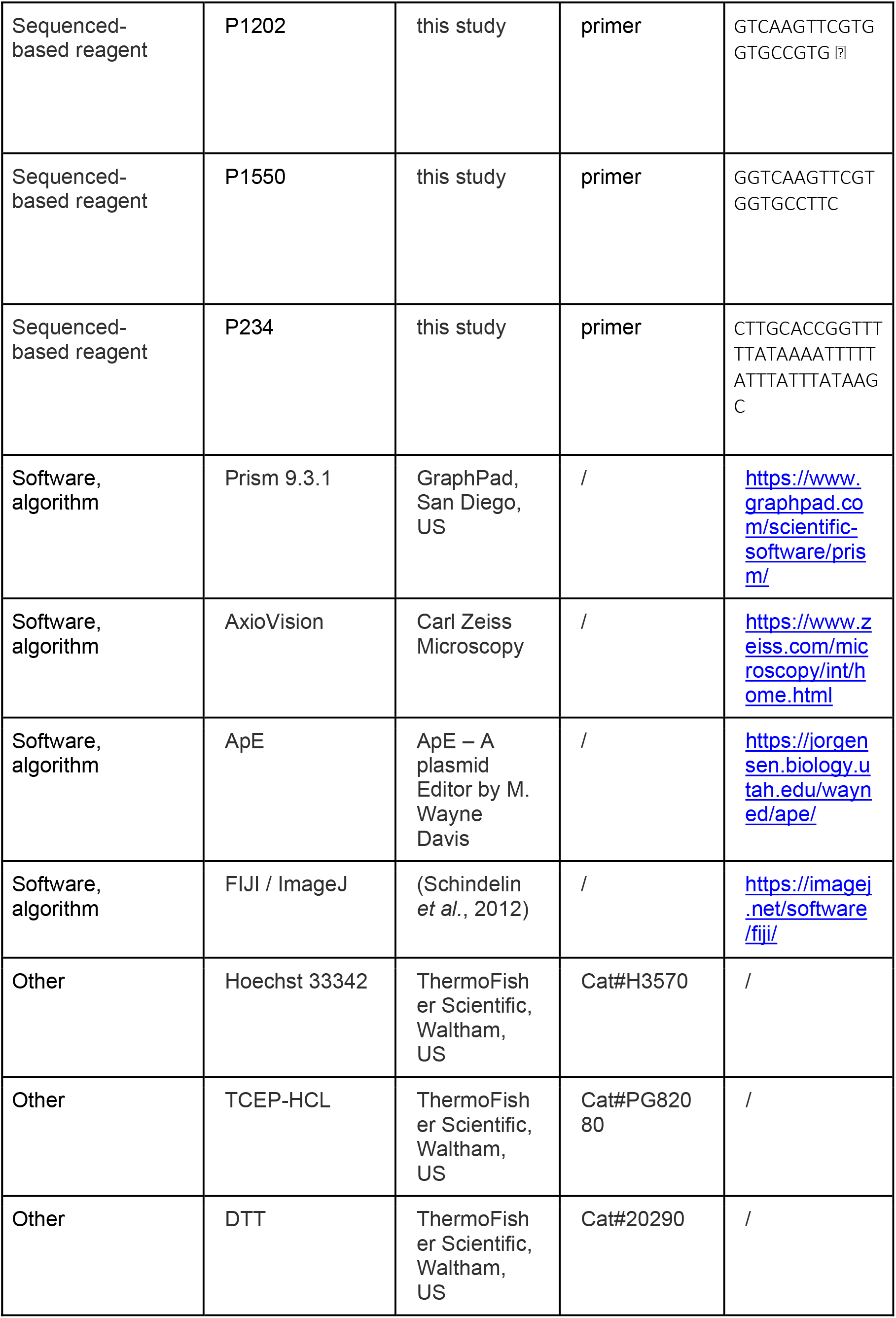

## Materials and Methods

### Animal models

Female 4–6-week-old Swiss CD1 or C57BL/6 mice from Janvier or Charles River laboratories were used to propagate *P. berghei* parasites. Transgenic parasites were generated on the *Plasmodium berghei* ANKA background (Vincke & Bafort, 1968) either directly in wild-type or wild-type-derived strains (e.g., *trap(−)* and *cfluo*). Parasites were cultured in Swiss CD1 mice, whereas transmission experiments with sporozoites were performed only in C57Bl/6 mice.

### Generation of *S210C*, *S210c/Q216C* and *S210C/F224C* parasites

To generate the parasite lines *S210C*, *S210C/Q216C* and *S210C/F224C* we made use of a synthetic *trap* gene that had been codon modified for *E. coli* K12 and used previously to create I domain exchange mutants (Klug et al., 2020). The synthetic *trap* gene cloned in the pMK-RQ vector (Invitrogen) was targeted by site-directed mutagenesis using the primers P1149/P1150 (S210C), P1153/P1154 (Q216C) and P1151/P1152 (F224C). S210C/Q216C and S210C/F224C mutations were introduced sequentially. Mutated sequences were cloned into the Pb238-TRAP-NdeI/PacI vector (*NdeI/PacI*), linearized (*ScaI-HF*) and transfected into *P. berghei* ANKA using standard protocols (Janse et al., 2006). Isogenic parasite populations were generated as described previously (Klug et al., 2020).

### Mosquito infection

Mice were infected intraperitoneally by injection of frozen parasite stabilates(~200-250 μl). Subsequently, infected mice were bled by cardiac puncture once parasitemia reached ~2% and used for a fresh blood transfer of 20.000.000 parasites into two naïve mice. Mice that obtained a blood transfer were kept for further 3-4 days. Mice were fed to mosquitoes if at least three exflagellation events per field of view were observed. Mice that contained the right density of gametocytes were anaesthetized with a mixture of ketamine and xylazine (87.5 mg/kg ketamine and 12.5 mg/kg xylazine), placed on mosquito cages and covered with paper tissues to dim the light and enhance mosquito biting. Mosquitoes were allowed to take a blood meal for 20-30 min on at least two mice per mosquito cage. Mosquitos were shaken off twice during this period to guarantee that most mosquitoes had the chance to suck blood. Subsequently infected mosquitoes were kept at 80% humidity and 21°C in a climate chamber. Mosquitoes determined to be infected had to be three to seven days old and were starved over night prior to blood feeding.

### Preparation of hemolymph, midgut and salivary gland sporozoites

Sporozoites were isolated from midguts, hemolymph and salivary glands of infected mosquitos between day 12 and day 22 post infection. The timepoint for dissection was dependent on the planned experiments; midgut sporozoites were dissected between day 12 and 15, hemolymph sporozoites between day 15 and 16 and salivary gland sporozoites between day 17 and 22 post infection. For counting experiments, midguts and salivary glands of at least 10 mosquitoes were dissected in PBS, the tissue was crushed with a pestle and free sporozoites were counted using a Neubauer counting chamber. The counting chamber was loaded with 10 μl solution from the side and sporozoites were allowed to settle for 5 min prior to counting. Sporozoites were counted using a light microscope (Carl Zeiss) and 400-fold magnification. To isolate hemolymph sporozoites, mosquitoes were anaesthetized by cooling on ice for at least 10 min. Once mosquitoes were immobile the last segment of the abdomen was cut with a syringe. Prepared mosquitoes were flushed by inserting a long drawn Pasteur pipette into the lateral side of the thorax and injected with RPMI (supplemented with 50.000 units/l penicillin and 50 mg/l streptomycin). The hemolymph was thus drained from the abdomen, collected on a piece of foil and transferred to a plastic reaction tube (Eppendorf). Hemolymph sporozoites were counted as previously described for midgut and salivary gland sporozoites.

### Gliding assays under regular and reducing conditions

To reduce introduced disulphide bonds fixing the TRAP I domain in either the closed or open confirmation a reducing agent was added prior to the gliding essay. HL sporozoites were dissected according to standard protocols. Samples were centrifuged for one min at 13.000 rpm (Thermo Fisher Scientific, Biofuge primo), supernatant was removed and replaced with DTT or TCEP diluted in RPMI at various concentrations (DTT: 50 mM - 1000 mM, TCEP: 5-200 mM) and incubated for varying lengths of time (1-30 min). Subsequently samples were centrifuged and sporozoite pellets were washed three times with RPMI. Finally, after the last washing step RPMI was replaced with RPMI containing 3% BSA to activate sporozoites and samples were transferred to a 96-well plate with optical bottom (Nunc). To allow sporozoites to attach, plates were centrifuged at 1500 rpm for three min (Heraeus Multifuge S1). Fluorescence microscopy (Axiovert 200M, Carl Zeiss) with a 25x objective was used to capture 1-3 minute movies with a frame rate of three seconds. The movies were analyzed using the Manual Tracking plugin of ImageJ (Schindelin et al., 2012) to determine the speed and trajectories of the moving sporozoites **(see Classification of Sporozoites)**. For standard sporozoite gliding assays the treament with TCEP or DTT as well as following washing steps were omitted. Instead sporozoites were directly activated with RPMI containing 3% BSA and imaged as described.

### Classification of sporozoite movement patterns

Movement is described as gliding if sporozoites travel at least one full circle in a 1 min time-lapse. Unproductive but motile forms of gliding are patch gliding and waving. Patch gliding describes sporozoites moving back and forth over a single adhesion site. In the maximum projection patch gliding sporozoites show a haystack pattern. Waving sporozoites are attached at a single adhesion side, swinging around with the other end. Non-motile sporozoites can either be attached or free floating in the medium. For visual examples please see **Figure S2**.

### Infection by mosquito bites and sporozoite injections

To determine the transmission potential of generated parasite lines mice were infected by mosquito bites and sporozoite injections. To study native transmission mosquitoes that had been infected 17-24 days before were screened for the presence of sporozoites visible as GFP positive midguts, separated in cups of 10 each and starved for 6–8 h. Subsequently naive female C57Bl/6 mice were anaesthetized by intraperitoneal injection of a mixture of ketamine and xylazine (87.5 mg/kg ketamine and 12.5 mg/kg xylazine). Subsequently anaesthetized mice were placed with the ventral side on the prepared cups for approximately 20 min. Mosquitoes that had taken a blood meal were dissected afterwards to randomly determine sporozoite numbers within salivary glands and midguts. For the injection of hemolymph sporozoites the hemolymph of mosquitoes that had been infected 13 to 16 days before was obtained as described previously. Sporozoites were counted in a Neubauer counting chamber and diluted with PBS to 10.000 hemolymph sporozoites per 100 μl. If sporozoite concentrations were too low, samples were centrifuged for 1 min at 10.000 rpm (Thermo Fisher Scientific, Biofuge primo). Subsequently the excess of liquid was removed and sporozoites were resuspended in an appropriate volume to achieve the planned dilution. For the injection of salivary gland sporozoites the salivary glands of mosquitoes that had been infected 17 to 24 days earlier were dissected in PBS. Sporozoites were released as described above and diluted with PBS to 10.000 salivary gland sporozoites per 100 μl. Sporozoite solutions were injected intravenously in the tail vein of naive C57Bl/6 mice. The parasitemia of infected mice was monitored by daily blood smears from day 3 on up to day 20 post infection. In addition, the survival of infected mice was monitored up to 20 days. Blood smears were stained in Giemsa solution (Merck) and counted using a light microscope (Carl Zeiss) with a counting grid. The time difference between the infection and the observation of the first blood stage was determined as prepatency.

### Antibodies

For immunofluorescence stainings and western blots we made use of antibodies directed against the circumsporozoite protein (CSP) and the thrombospondin related anonymous protein (TRAP). The anti-CSP antibody mAb 3D11 (Yoshida et al., 1980) was applied as unpurified culture supernatant of the corresponding hybridoma cell line (1:200 diluted for immunofluorescence assays and 1:1.000 diluted for western blots). TRAP antibodies were generated as described previously (Klug et al., 2020) and applied as 1:200 dilution in immunofluorescence stainings and as 1:1.000 dilution for western blots. Secondary antibodies coupled to AlexaFluor 488 (Goat anti-Rabbit), Atto647N (Goat anti-Rabbit) and Atto 594 (Goat anti-Mouse) were obtained from Thermo-Fisher and Sigma-Aldrich and used as 1:200 dilution in immunofluorescence stainings to visualize localization of TRAP and CSP. To avoid spillover of fluorescence signals in *P. berghei* lines expressing mCherry at the sporozoite stage Atto647N secondary antibodies were applied to visualize TRAP while CSP and TRAP in non-fluorescent *P. berghei* lines were stained using AlexaFluor488 and Atto594 secondary antibodies. For western blots secondary antibodies coupled to HRP (Bio-Rad and GE-Healthcare) at 1:10.000 dilution were applied (Sheep anti-Mouse, Goat anti-Rabbit).

### Immunofluorescence staining

Localization of TRAP and CSP was visualized by immunofluorescence staining of hemolymph sporozoites. Hemolymph was isolated from *P. berghei* infected mosquitoes as described previously (Klug et al., 2020) and collected in plastic reaction tubes. Obtained hemolymph sporozoites were transferred into an 8-well glass-bottom imaging chamber (Nunc Lab-Tek), activated with 3 % BSA in RPMI medium and forced to attach to the bottom by centrifugation at 800rpm for 8 min. Subsequently sporozoites were allowed to glide for 15 min at RT. Afterwards samples were fixated using 4% PFA (in PBS) for 1h at RT. Fixed samples were washed three times with PBS and treated with primary antibody solutions for 1h at RT. Subsequently samples were washed three times with PBS and treated with secondary antibody solutions for 1h at RT in the dark. Finally, samples were washed three times in PBS and the supernatant was discarded. Samples were examined directly using a spinning disc confocal microscope (Nikon Ti series) with 100-fold magnification (Plan Apo VC 100x/1.4 N.A. oil immersion) and a Hamamatsu sCMOS ORCA Flash 4.0 camera.

### Western blotting

Western blotting was performed as described in (Klug et al., 2020). Hemolymph sporozoites were isolated from infected mosquitoes by flushing with PBS using a drawn out pipette as described before. Sporozoite solutions were kept on ice, counted using a hemocytometer and distributed to 30.000 sporozoites per reaction tube. Samples were centrifuged for 1 min with 13.000 rpm (Thermo Fisher Scientific, Biofuge primo) at 4°C to pellet sporozoites. Subsequently the supernatant was discarded, and pellets were lysed in 20 μL RIPA buffer (50 mM Tris pH 8, 1% NP40, 0.5% sodium dexoycholate, 0.1% SDS, 150 mM NaCl, 2 mM EDTA) supplemented with protease inhibitors (Sigma-Aldrich, P8340) for ≥1 hr on ice. Lysates were mixed with Laemmli buffer, heated for 10 min at 95°C and centrifuged for 1 min at 13.000 rpm (Thermo Fisher Scientific, Biofuge primo). Samples were separated on precast 4–15% SDS-PAGE gels (Mini Protein TGX Gels, Bio-Rad) and blotted on nitrocellulose membranes with the Trans-Blot Turbo Transfer System (Bio-Rad). Blocking was performed by incubation in PBS containing 0.05% Tween20 and 5% milk powder for 1 hr at RT. Afterwards, the solution was refreshed and antibodies directed against TRAP (rabbit polyclonal antibody, 1:1000 diluted) or the loading control CSP (mAb 3D11, cell culture supernatant 1:1.000 diluted) were added. Membranes were washed three times (PBS with 0.05% Tween20) and secondary anti-rabbit antibodies (Immun-Star (GAR)-HRP, Bio-Rad) or anti-mouse antibodies (NXA931, GE Healthcare) conjugated to horse radish peroxidase were applied for 1 hr (1:10.000 dilution) at room temperature. Signals were detected using SuperSignal West Pico Chemiluminescent Substrate or SuperSignal West Femto Maximum Sensitivity Substrate (Thermo Fisher Scientific). After the detection of TRAP, blots were treated with stripping buffer (Glycine 15 g/L, SDS 1 g/L, Tween20 10 ml/L, pH 2,2) for 15 min prior to incubation with anti-CSP antibodies used as loading control.

### Statistical analysis

Statistical analysis was performed using GraphPad Prism 9.3.1 (GraphPad, San Diego, CA, USA). Data sets were either tested with a one-way ANOVA (Kruskal-Wallis-test) or a Student’s t test. A value of p<0.05 was considered significant.

## Supplementary figures

**Figure S1:**
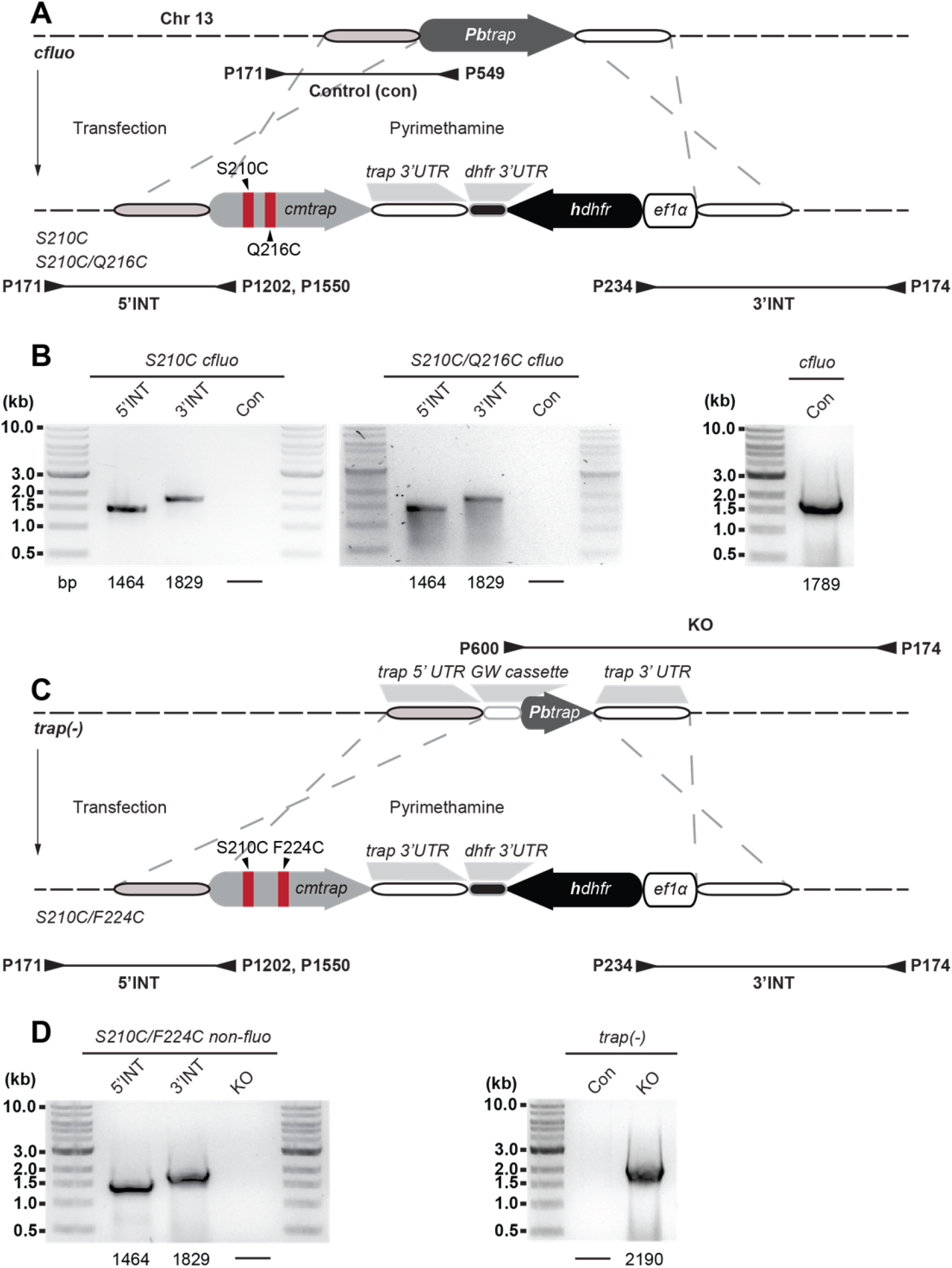
Generation of the recombinant parasite lines. **A, C**: TRAP genes with mutated I domains were transfected in *cFluo* or *trap(−)* parasites. Three different parasite lines were generated: *S210C*; control line with only one introduced cysteine, *S210C/Q216C*; mutant with two introduced cysteines which form a disulfide bond that fixates the I domain in the „open“ conformation and *S210C/F224C*; mutant with two introduced cysteines which form a disulfide bond that fixates the I domain in the „closed“ conformation. The TRAP gene in all three generated lines was codon modified for *E. coli* K12. Binding sites of primers used for genotyping and lengths of PCR products are indicated below the scheme. Note that the illustration is not drawn to scale. **B, D**: To control for correct integration of the transfected DNA sequences three different PCRs were performed amplifying sequences that are specific for successful DNA integration at the 5’ (5’INT) and the 3’ (3’INT) end as well as a PCR that is specific for the recipient line (Con/Control or Knockout/KO). The lengths of the expected PCR products are depicted below the images. Shown are only PCR results of isogenic populations cloned by limiting dilution. To ensure the correct replacement of the native TRAP gene the integration site of transfected parasites was sequenced.

**Figure S2:**
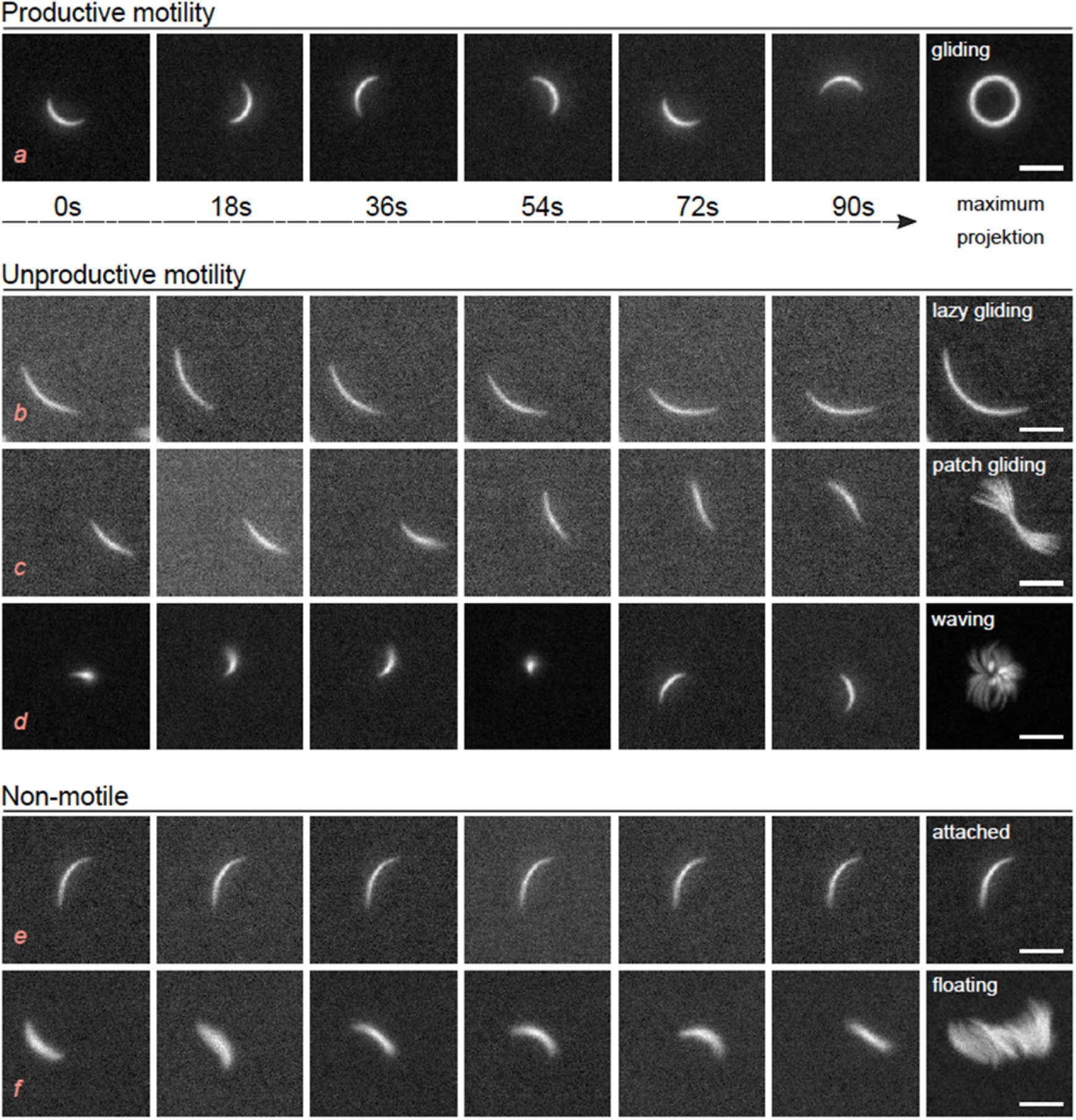
Movement patterns of *P. berghei* sporozoites. Observed sporozoite movement patterns from movies with stills taken every 18 seconds. Right image shows maximum fluorescence projection. Active movement can be divided in productive gliding motility (top), and unproductive movements such as lazy gliding for sporozoites gliding extremely slow, patch gliding for sporozoites moving rapidly over a single adhesion site in a back-and-forth manner, waving for sporozoites attached with one end and moving the rest of the sporozoite body within the medium. Non-motile sporozoites can be either attached to the substrate or floating in the medium. Scale bars: 5 μm. Red italic lettering is used to make reference to the movement patterns in S3.

**Figure S3:**
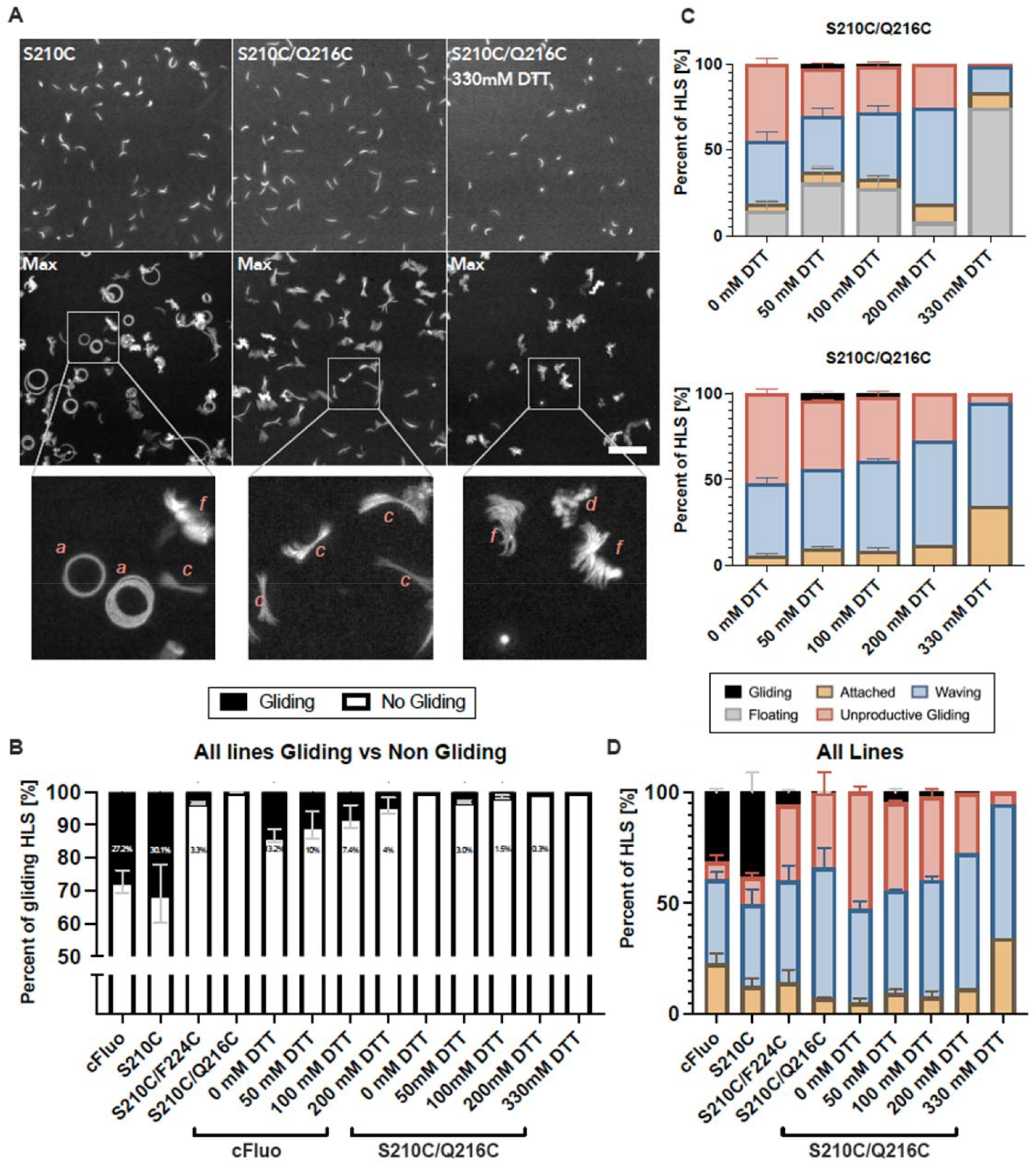
Reducing disulphide bonds with DTT rescues motility of sporozoites expressing the open I domain. **A**: Sporozoites do not show movement at high concentrations of DTT. Top: still images of hemolymph sporozoites of the S210C line, S210C/Q216C line, and S210C/Q216C line treated with 330 mM DTT. Bottom: Maximum projections of movies for the same lines. Circular movement pattern indicate productively moving sporozoites. For the open conformation, patch gliding sporozoites (haystack pattern) can be observed if no DTT is added. At a concentration of 330 mM DTT only floating sporozoites can be observed. In the bottom row, zoom-ins show sample sporozoites categorized into different movement patterns according to S2 (red lettering). Scale bar: 30μm. **B**: DTT rescues productive gliding in the open conformation. Gliding vs. non gliding hemolymph sporozoites (HLS) of all lines. Floaters have been included for all lines. **C**: Analysis of movement pattern of hemolymph derived sporozoites (HLS) of the indicated parasite lines with and without floaters. See Figure S2 for the classification of different movement patterns. Floaters have been omitted for the lower graph. **D**: Movement type analysis of hemolymph derived sporozoites of the indicated parasite lines without floaters. Shown results represent the same datasets as in (B) and (C) in a combined manner and broken down by movement patterns. Note that sporozoites treated with DTT (including the 0 mM DTT control) have gone through an additional incubation and washing step. See Figure S2 for the classification of different movement patterns.

## Bibliography

Aliprandini, E., Tavares, J., Panatieri, R. H., Thiberge, S., Yamamoto, M. M., Silvie, O., Ishino, T., Yuda, M., Dartevelle, S., Traincard, F., Boscardin, S. B., & Amino, R. (2018). Cytotoxic anti-circumsporozoite antibodies target malaria sporozoites in the host skin. Nat Microbiol, 3(11), 1224–1233. https://doi.org/10.1038/s41564-018-0254-z

Bane, K. S., Lepper, S., Kehrer, J., Sattler, J. M., Singer, M., Reinig, M., Klug, D., Heiss, K., Baum, J., Mueller, A. K., & Frischknecht, F. (2016). The Actin Filament-Binding Protein Coronin Regulates Motility in Plasmodium Sporozoites. PLoS Pathog, 12(7), e1005710. https://doi.org/10.1371/journal.ppat.1005710

Carey, A. F., Singer, M., Bargieri, D., Thiberge, S., Frischknecht, F., Ménard, R., & Amino, R. (2014). Calcium dynamics of Plasmodium berghei sporozoite motility. Cell Microbiol, 16(5), 768–783. https://doi.org/10.1111/cmi.12289

Cline, D. J., Redding, S. E., Brohawn, S. G., Psathas, J. N., Schneider, J. P., & Thorpe, C. (2004). New water-soluble phosphines as reductants of peptide and protein disulfide bonds: reactivity and membrane permeability. Biochemistry, 43(48), 15195–15203. https://doi.org/10.1021/bi048329a

Coppi, A., Natarajan, R., Pradel, G., Bennett, B. L., James, E. R., Roggero, M. A., Corradin, G., Persson, C., Tewari, R., & Sinnis, P. (2011). The malaria circumsporozoite protein has two functional domains, each with distinct roles as sporozoites journey from mosquito to mammalian host. J Exp Med, 208(2), 341–356. https://doi.org/10.1084/jem.20101488

Doud, M. B., Koksal, A. C., Mi, L. Z., Song, G., Lu, C., & Springer, T. A. (2012). Unexpected fold in the circumsporozoite protein target of malaria vaccines. Proc Natl Acad Sci U S A, 109(20), 7817–7822. https://doi.org/10.1073/pnas.1205737109

Douglas, R. G., Amino, R., Sinnis, P., & Frischknecht, F. (2015). Active migration and passive transport of malaria parasites. Trends Parasitol, 31(8), 357–362. https://doi.org/10.1016/j.pt.2015.04.010

Dundas, K., Shears, M. J., Sun, Y., Hopp, C. S., Crosnier, C., Metcalf, T., Girling, G., Sinnis, P., Billker, O., & Wright, G. J. (2018). Alpha-v-containing integrins are host receptors for the Plasmodium falciparum sporozoite surface protein, TRAP. Proc Natl Acad Sci U S A, 115(17), 4477–4482. https://doi.org/10.1073/pnas.1719660115

Ejigiri, I., Ragheb, D. R., Pino, P., Coppi, A., Bennett, B. L., Soldati-Favre, D., & Sinnis, P. (2012). Shedding of TRAP by a rhomboid protease from the malaria sporozoite surface is essential for gliding motility and sporozoite infectivity. PLoS Pathog, 8(7), e1002725. https://doi.org/10.1371/journal.ppat.1002725

Flores-Garcia, Y., Wang, L. T., Park, M., Asady, B., Idris, A. H., Kisalu, N. K., Munoz, C., Pereira, L. S., Francica, J. R., Seder, R. A., & Zavala, F. (2021). The P. falciparum CSP repeat region contains three distinct epitopes required for protection by antibodies in vivo. PLoS Pathog, 17(11), e1010042. https://doi.org/10.1371/journal.ppat.1010042

Frischknecht, F., & Matuschewski, K. (2017). Plasmodium Sporozoite Biology. Cold Spring Harb Perspect Med, 7(5). https://doi.org/10.1101/cshperspect.a025478

Gantt, S., Persson, C., Rose, K., Birkett, A. J., Abagyan, R., & Nussenzweig, V. (2000). Antibodies against thrombospondin-related anonymous protein do not inhibit Plasmodium sporozoite infectivity in vivo. Infect Immun, 68(6), 3667–3673. http://www.ncbi.nlm.nih.gov/entrez/query.fcgi?cmd=Retrieve&db=PubMed&dopt=Citation&list_uids=10816526

Gaudinski, M. R., Berkowitz, N. M., Idris, A. H., Coates, E. E., Holman, L. A., Mendoza, F., Gordon, I. J., Plummer, S. H., Trofymenko, O., Hu, Z., Campos Chagas, A., O’Connell, S., Basappa, M., Douek, N., Narpala, S. R., Barry, C. R., Widge, A. T., Hicks, R., Awan, S. F., … Team, V. R. C. S. (2021). A Monoclonal Antibody for Malaria Prevention. N Engl J Med, 385(9), 803–814. https://doi.org/10.1056/NEJMoa2034031

Ghosh, A. K., Devenport, M., Jethwaney, D., Kalume, D. E., Pandey, A., Anderson, V. E., Sultan, A. A., Kumar, N., & Jacobs-Lorena, M. (2009). Malaria parasite invasion of the mosquito salivary gland requires interaction between the Plasmodium TRAP and the Anopheles saglin proteins. PLoS Pathog, 5(1), e1000265. https://doi.org/10.1371/journal.ppat.1000265

Heintzelman, M. B. (2015). Gliding motility in apicomplexan parasites. Semin Cell Dev Biol, 46, 135–142. https://doi.org/10.1016/j.semcdb.2015.09.020

Herrera, R., Anderson, C., Kumar, K., Molina-Cruz, A., Nguyen, V., Burkhardt, M., Reiter, K., Shimp, R., Jr., Howard, R. F., Srinivasan, P., Nold, M. J., Ragheb, D., Shi, L., DeCotiis, M., Aebig, J., Lambert, L., Rausch, K. M., Muratova, O., Jin, A., … Narum, D. L. (2015). Reversible Conformational Change in the Plasmodium falciparum Circumsporozoite Protein Masks Its Adhesion Domains. Infect Immun, 83(10), 3771–3780. https://doi.org/10.1128/IAI.02676-14

Hopp, C. S., Kanatani, S., Archer, N. K., Miller, R. J., Liu, H., Chiou, K. K., Miller, L. S., & Sinnis, P. (2021). Comparative intravital imaging of human and rodent malaria sporozoites reveals the skin is not a species-specific barrier. EMBO Mol Med, 13(4), e11796. https://doi.org/10.15252/emmm.201911796

Janse, C. J., Franke-Fayard, B., Mair, G. R., Ramesar, J., Thiel, C., Engelmann, S., Matuschewski, K., van Gemert, G. J., Sauerwein, R. W., & Waters, A. P. (2006). High efficiency transfection of Plasmodium berghei facilitates novel selection procedures. Mol Biochem Parasitol, 145(1), 60–70. https://doi.org/10.1016/j.molbiopara.2005.09.007

Jethwaney, D., Lepore, T., Hassan, S., Mello, K., Rangarajan, R., Jahnen-Dechent, W., Wirth, D., & Sultan, A. A. (2005). Fetuin-A, a hepatocyte-specific protein that binds Plasmodium berghei thrombospondin-related adhesive protein: a potential role in infectivity. Infect Immun, 73(9), 5883–5891. https://doi.org/10.1128/IAI.73.9.5883-5891.2005

Julien, J. P., & Wardemann, H. (2019). Antibodies against Plasmodium falciparum malaria at the molecular level. Nat Rev Immunol, 19(12), 761–775. https://doi.org/10.1038/s41577-019-0209-5

Kehrer, J., Singer, M., Lemgruber, L., Silva, P. A., Frischknecht, F., & Mair, G. R. (2016). A Putative Small Solute Transporter Is Responsible for the Secretion of G377 and TRAP-Containing Secretory Vesicles during Plasmodium Gamete Egress and Sporozoite Motility. PLoS Pathog, 12(7), e1005734. https://doi.org/10.1371/journal.ppat.1005734

Kim, Y. C., Dema, B., Rodriguez-Garcia, R., Lopez-Camacho, C., Leoratti, F. M. S., Lall, A., Remarque, E. J., Kocken, C. H. M., & Reyes-Sandoval, A. (2020). Evaluation of Chimpanzee Adenovirus and MVA Expressing TRAP and CSP from Plasmodium cynomolgi to Prevent Malaria Relapse in Nonhuman Primates. Vaccines (Basel), 8(3). https://doi.org/10.3390/vaccines8030363

Klug, D., Goellner, S., Kehrer, J., Sattler, J., Strauss, L., Singer, M., Lu, C., Springer, T. A., & Frischknecht, F. (2020). Evolutionarily distant I domains can functionally replace the essential ligand-binding domain of Plasmodium TRAP. Elife, 9. https://doi.org/10.7554/eLife.57572

Li, J., & Springer, T. A. (2017). Integrin extension enables ultrasensitive regulation by cytoskeletal force. Proc Natl Acad Sci U S A, 114(18), 4685–4690. https://doi.org/10.1073/pnas.1704171114

Lu, C., Song, G., Beale, K., Yan, J., Garst, E., Feng, J., Lund, E., Catteruccia, F., & Springer, T. A. (2020). Design and assessment of TRAP-CSP fusion antigens as effective malaria vaccines. PLoS One, 15(1), e0216260. https://doi.org/10.1371/journal.pone.0216260

Matuschewski, K., Nunes, A. C., Nussenzweig, V., & Menard, R. (2002). Plasmodium sporozoite invasion into insect and mammalian cells is directed by the same dual binding system. EMBO J, 21(7), 1597–1606. https://doi.org/10.1093/emboj/21.7.1597

Menard, R., Tavares, J., Cockburn, I., Markus, M., Zavala, F., & Amino, R. (2013). Looking under the skin: the first steps in malarial infection and immunity. Nat Rev Microbiol, 11(10), 701–712. https://doi.org/10.1038/nrmicro3111

Münter, S., Sabass, B., Selhuber-Unkel, C., Kudryashev, M., Hegge, S., Engel, U., Spatz, J. P., Matuschewski, K., Schwarz, U. S., & Frischknecht, F. (2009). Plasmodium sporozoite motility is modulated by the turnover of discrete adhesion sites. Cell Host Microbe, 6(6), 551–562. https://doi.org/10.1016/j.chom.2009.11.007

Murugan, R., Scally, S. W., Costa, G., Mustafa, G., Thai, E., Decker, T., Bosch, A., Prieto, K., Levashina, E. A., Julien, J. P., & Wardemann, H. (2020). Evolution of protective human antibodies against Plasmodium falciparum circumsporozoite protein repeat motifs. Nat Med, 26(7), 1135–1145. https://doi.org/10.1038/s41591-020-0881-9

Nazeri, S., Zakeri, S., Mehrizi, A. A., Sardari, S., & Djadid, N. D. (2020). Measuring of IgG2c isotype instead of IgG2a in immunized C57BL/6 mice with Plasmodium vivax TRAP as a subunit vaccine candidate in order to correct interpretation of Th1 versus Th2 immune response. Exp Parasitol, 216, 107944. https://doi.org/10.1016/j.exppara.2020.107944

Prudencio, M., Rodriguez, A., & Mota, M. M. (2006). The silent path to thousands of merozoites: the Plasmodium liver stage. Nat Rev Microbiol, 4(11), 849–856. https://doi.org/10.1038/nrmicro1529

Quadt, K. A., Streichfuss, M., Moreau, C. A., Spatz, J. P., & Frischknecht, F. (2016). Coupling of Retrograde Flow to Force Production During Malaria Parasite Migration. ACS Nano, 10(2), 2091–2102. https://doi.org/10.1021/acsnano.5b06417

Schindelin, J., Arganda-Carreras, I., Frise, E., Kaynig, V., Longair, M., Pietzsch, T., Preibisch, S., Rueden, C., Saalfeld, S., Schmid, B., Tinevez, J. Y., White, D. J., Hartenstein, V., Eliceiri, K., Tomancak, P., & Cardona, A. (2012). Fiji: an open-source platform for biological-image analysis. Nat Methods, 9(7), 676–682. https://doi.org/10.1038/nmeth.2019

Shimaoka, M., Lu, C., Palframan, R. T., von Andrian, U. H., McCormack, A., Takagi, J., & Springer, T. A. (2001). Reversibly locking a protein fold in an active conformation with a disulfide bond: integrin alphaL I domains with high affinity and antagonist activity in vivo. Proc Natl Acad Sci U S A, 98(11), 6009–6014. https://doi.org/10.1073/pnas.101130498

Shimaoka, M., Takagi, J., & Springer, T. A. (2002). Conformational regulation of integrin structure and function. Annu Rev Biophys Biomol Struct, 31, 485–516. https://doi.org/10.1146/annurev.biophys.31.101101.140922

Shimaoka, M., Xiao, T., Liu, J. H., Yang, Y., Dong, Y., Jun, C. D., McCormack, A., Zhang, R., Joachimiak, A., Takagi, J., Wang, J. H., & Springer, T. A. (2003). Structures of the alpha L I domain and its complex with ICAM-1 reveal a shape-shifting pathway for integrin regulation. Cell, 112(1), 99–111. https://doi.org/10.1016/s0092-8674(02)01257-6

Song, G., Koksal, A. C., Lu, C., & Springer, T. A. (2012). Shape change in the receptor for gliding motility in Plasmodium sporozoites. Proc Natl Acad Sci U S A, 109(52), 21420–21425. https://doi.org/10.1073/pnas.1218581109

Spaccapelo, R., Naitza, S., Robson, K. J., & Crisanti, A. (1997). Thrombospondin-related adhesive protein (TRAP) of Plasmodium berghei and parasite motility. Lancet, 350(9074), 335. https://doi.org/10.1016/S0140-6736(97)24031-6

Springer, T. A. (2006). Complement and the multifaceted functions of VWA and integrin I domains. Structure, 14(11), 1611–1616. https://doi.org/10.1016/j.str.2006.10.001

Steel, R. W. J., Vigdorovich, V., Dambrauskas, N., Wilder, B. K., Arredondo, S. A., Goswami, D., Kumar, S., Carbonetti, S., Swearingen, K. E., Nguyen, T., Betz, W., Camargo, N., Fisher, B. S., Soden, J., Thomas, H., Freeth, J., Moritz, R. L., Noah Sather, D., & Kappe, S. H. I. (2021). Platelet derived growth factor receptor beta (PDGFRbeta) is a host receptor for the human malaria parasite adhesin TRAP. Sci Rep, 11(1), 11328. https://doi.org/10.1038/s41598-021-90722-5

Sultan, A. A., Thathy, V., Frevert, U., Robson, K. J., Crisanti, A., Nussenzweig, V., Nussenzweig, R. S., & Menard, R. (1997). TRAP is necessary for gliding motility and infectivity of plasmodium sporozoites. Cell, 90(3), 511–522. https://doi.org/10.1016/s0092-8674(00)80511-5

Tiono, A. B., Nebie, I., Anagnostou, N., Coulibaly, A. S., Bowyer, G., Lam, E., Bougouma, E. C., Ouedraogo, A., Yaro, J. B. B., Barry, A., Roberts, R., Rampling, T., Bliss, C., Hodgson, S., Lawrie, A., Ouedraogo, A., Imoukhuede, E. B., Ewer, K. J., Viebig, N. K., Sirima, S. B. (2018). First field efficacy trial of the ChAd63 MVA ME-TRAP vectored malaria vaccine candidate in 5-17 months old infants and children. PLoS One, 13(12), e0208328. https://doi.org/10.1371/journal.pone.0208328

Vanderberg, J. P. (1974). Studies on the motility of Plasmodium sporozoites. J Protozool, 21(4), 527–537. https://doi.org/10.1111/j.1550-7408.1974.tb03693.x

Vincke, I. H., & Bafort, J. (1968). [Standardization of the inoculum of Plasmodium berghei sporozoites]. Ann Soc Belges Med Trop Parasitol Mycol, 48(2), 181–194. https://www.ncbi.nlm.nih.gov/pubmed/5744433 (Methodes de standardisation de l’inoculum de sporozoites de Plasmodium berghei.)

Wang, L. T., Pereira, L. S., Kiyuka, P. K., Schon, A., Kisalu, N. K., Vistein, R., Dillon, M., Bonilla, B. G., Molina-Cruz, A., Barillas-Mury, C., Tan, J., Idris, A. H., Francica, J. R., & Seder, R. A. (2021). Protective effects of combining monoclonal antibodies and vaccines against the Plasmodium falciparum circumsporozoite protein. PLoS Pathog, 17(12), e1010133. https://doi.org/10.1371/journal.ppat.1010133

Yang, M., Dutta, C., & Tiwari, A. (2015). Disulfide-bond scrambling promotes amorphous aggregates in lysozyme and bovine serum albumin. J Phys Chem B, 119(10), 3969–3981. https://doi.org/10.1021/acs.jpcb.5b00144

Yoshida, N., Nussenzweig, R. S., Potocnjak, P., Nussenzweig, V., & Aikawa, M. (1980). Hybridoma produces protective antibodies directed against the sporozoite stage of malaria parasite. Science, 207(4426), 71–73. https://doi.org/10.1126/science.6985745

